# MSH2 is not required for either maintenance of DNA methylation or repeat contraction at the *FMR1* locus in fragile X syndrome

**DOI:** 10.1101/2024.12.20.629815

**Authors:** Jessalyn Grant-Bier, Kathryn Ruppert, Bruce Hayward, Karen Usdin, Daman Kumari

## Abstract

**Background:** Repeat-induced epigenetic changes are observed in many repeat expansion disorders (REDs). These changes result in transcriptional deficits and/or silencing of the associated gene. MSH2, a mismatch repair protein that is required for repeat expansion in the REDs, has been implicated in the maintenance of DNA methylation seen in the region surrounding expanded CTG repeats at the *DMPK* locus in myotonic dystrophy type 1 (DM1). Here, we investigated the role of MSH2 in aberrant DNA methylation in two additional REDs, fragile X syndrome (FXS) that is caused by a CGG repeat expansion in the 5’ untranslated region (UTR) of the fragile X messenger ribonucleoprotein 1 (*FMR1*) gene, and Friedreich’s ataxia (FRDA) that is caused by GAA repeat expansion in intron 1 of the *frataxin* (*FXN*) gene.

**Results:** In contrast to what is seen at the *DMPK* locus in DM1, loss of MSH2 did not decrease DNA methylation at the *FMR1* promoter in FXS embryonic stem cells (ESCs) or increase *FMR1* transcription. This difference was not due to the differences in the CpG density of the two loci as a decrease in DNA methylation was also not observed in a less CpG dense region upstream of the expanded GAA repeats in the *FXN* gene in MSH2 null induced pluripotent stem cells (iPSCs) derived from FRDA patient fibroblasts. Surprisingly given previous reports, we found that *FMR1* reactivation was associated with a high frequency of MSH2- independent repeat contractions that resulted a permanent loss of DNA methylation.

**Conclusions:** Our results suggest that there are mechanistic differences in the way that DNA methylation is maintained in the vicinity of expanded repeats among different REDs even though they share a similar mechanism of repeat expansion. The high frequency of transcription-induced MSH2-independent contractions we have observed may contribute to the mosaicism that is frequently seen in carriers of *FMR1* alleles with expanded CGG-repeat tracts. Given the recent interest in the therapeutic use of transcription-driven repeat contractions, our data may have interesting mechanistic, prognostic, and therapeutic implications.

**Graphical abstract:** 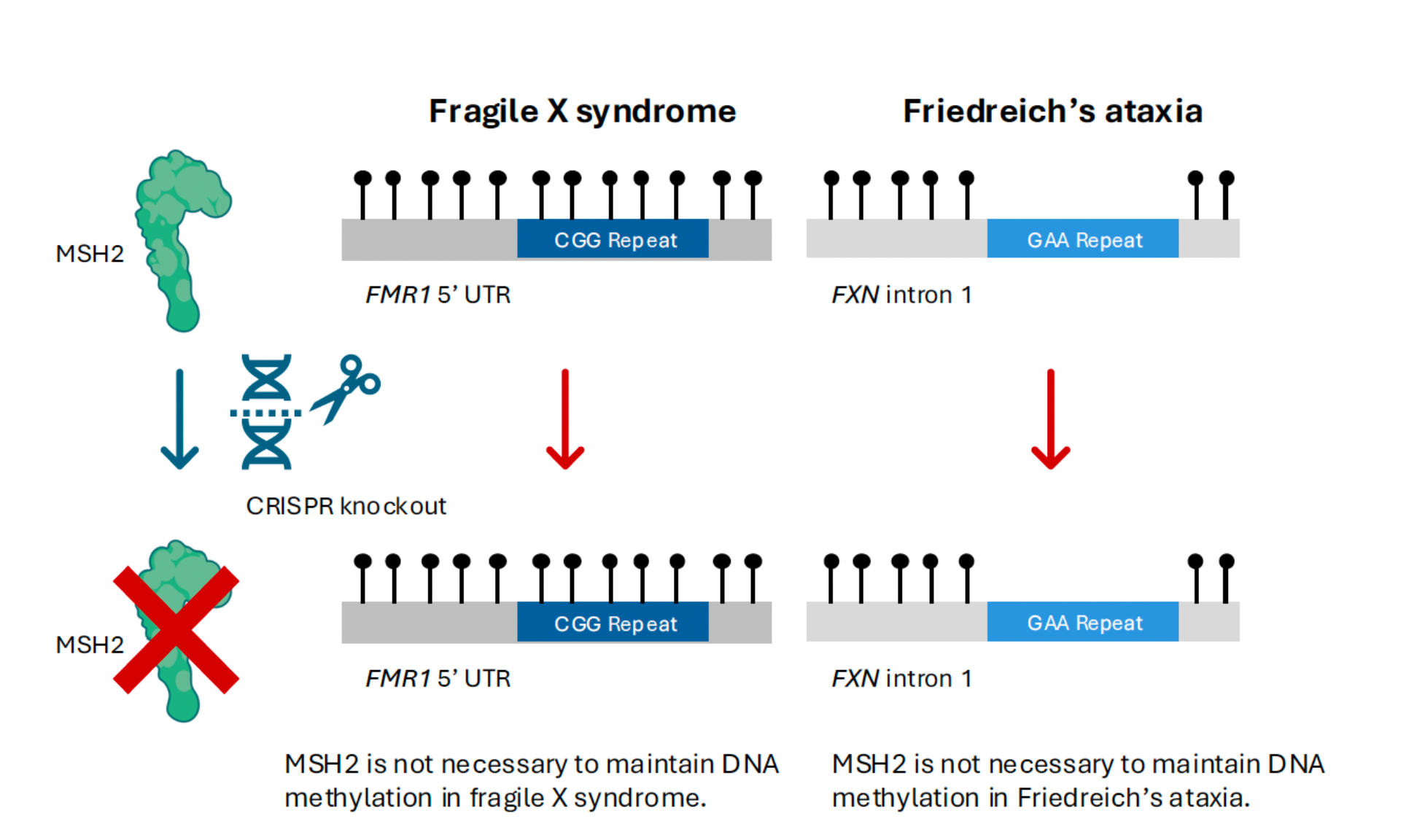

## Background

There are more than 50 human genetic disorders that are all caused by the expansion of a short tandem repeat tract in a single specific gene (1). Collectively, these are known as repeat expansion disorders (REDs). While the unit length, sequence and location of the repeat tract differ, all REDs share some common features. The expanded repeats form a variety of secondary structures (2, 3) that block DNA replication (4, 5, 6, 7, 8) and transcription (9, 10, 11, 12, 13). Such structures are thought to contribute to the instability of the repeats as well as to some of the downstream consequences of repeat expansion including epigenetic changes that negatively affect transcription.

Fragile X syndrome (FXS; OMIM #300624), an X-linked neurodevelopmental disorder, is caused by the expansion of a CGG trinucleotide repeat in the 5’ untranslated region (UTR) of the *Fragile X Messenger Ribonucleoprotein 1* (*FMR1)* gene (14). Typical *FMR1* alleles in unaffected individuals carry 6-54 CGG repeats. When the CGG repeat expands beyond a threshold of 200 repeats, the 5’ end of the gene including the promoter and the repeats themselves, become hypermethylated. This results in transcriptional silencing (15, 16) and the absence of FMRP, an RNA binding protein important for synaptic plasticity (17). Repeat expansion-induced DNA methylation has also been reported in other REDs including Friedreich’s ataxia (FRDA) and myotonic dystrophy type 1 (DM1). FRDA (OMIM #229300) is an autosomal recessive progressive neurodegenerative disorder caused by the expansion of a GAA repeat in intron 1 of the *frataxin* (*FXN*) gene. Typical *FXN* alleles in unaffected individuals have between 7-33 GAA repeats and FRDA patients have between 66-1700 repeats (18, 19). Repeat expansion causes a deficit of *FXN* mRNA by affecting both transcription initiation and elongation (10, 11). This results in reduced expression of frataxin, a ubiquitous and highly conserved mitochondrial protein (20). Some of the CpG residues surrounding the expanded GAA repeats are hypermethylated in FRDA cells (21, 22, 23). Increased methylation of CpG residues is also seen upstream of the expanded repeats in DM1 (OMIM #160900), an autosomal dominant disorder characterized by a CTG repeat expansion in the 3’ UTR of the *Dystrophy Myotonic Protein Kinase* (*DMPK)* gene (24, 25). A typical *DMPK* allele has 5-37 CTG repeats whereas DM1 patients have between 50- 5000 CTG repeats. The CTG repeat expansion affects the expression of *DMPK* and neighboring genes, *Sine Oculis Homeobox Homolog 5* (*SIX5*) and *Dystrophia Myotonica WD Repeat-Containing* (*DMWD*) and may contribute to the DM1 phenotype (26, 27, 28, 29). In addition to increased DNA methylation, *FMR1*, *FXN* and *DMPK* alleles with expanded repeats are all enriched for repressive histone modifications including histone H3 di- and tri-methylated at lysine 9 (22, 30, 31, 32, 33). A better understanding of why and how repeat expansion causes these epigenetic changes may allow the development of interventions that could restore gene expression in affected REDs.

MutS homolog 2 (MSH2) is a protein that dimerizes with either MSH6 or MSH3 to recognize and repair base mismatches and short insertion/deletion loops during MMR (34). Aside from its well-established role in repeat expansion (35, 36, 37, 38), MSH2 has recently been shown to play a role in repeat-mediated DNA methylation in DM1 (39). Loss of MSH2 resulted in reduced DNA methylation at CpG residues upstream of the CTG repeat in the *DMPK* gene in DM1 ESCs. Interestingly, this was also associated with contractions of the repeat. Reconstitution of MSH2 expression in null cell lines with large CTG repeats, that were above the threshold for DNA methylation, restored repeat expansions but did not re- methylate the region, suggesting a role for MSH2 in maintenance rather than *de novo* methylation in DM1 (39).

A similar role for MSH2 in the maintenance of gene silencing has been reported for *DUX4*, a developmentally silenced sub-telomeric gene, in HEK293T cells (40). MSH2 has also been shown to recruit DNA methyltransferase 1 (DNMT1) to sites of oxidative damage (41). The resulting DNA methylation has been suggested to cause a transient decrease in transcription to limit damage during the repair process, and the epigenetic state of the region is expected to be restored once DNA repair is completed (41). However, persistent DNA damage has been shown to result in heritable gene silencing (42). Loci with expanded CGG, GAA and CTG repeats are associated with increased DNA damage that causes repeat instability, and are prone to replication stress that has been shown to result in chromosome fragility (4, 7, 43, 44, 45, 46, 47). Moreover, stable R-loops are also formed at transcriptionally active *FMR1* and *FXN* alleles (48, 49, 50), which could also lead to increased DNA damage at both loci (51, 52). Given the propensity for increased DNA damage, it was possible that MSH2 also plays a role in aberrant DNA methylation at *FMR1* and *FXN* alleles with expanded repeats.

To test this hypothesis, we used the CRISPR/Cas9 system to generate MSH2 null cell lines from FXS ESCs with 400 CGG repeats in the *FMR1* gene and from induced pluripotent stem cells (iPSCs) derived from FRDA fibroblasts with 700-800 GAA repeats in the *FXN* gene. We studied the role of MSH2 in maintaining DNA methylation in the CpG dense *FMR1* gene promoter and the less-dense CpG region upstream of the GAA repeats in intron 1 of the *FXN* gene. We observed no difference in the CpG methylation in MSH2 null cell lines from both FXS ESCs and FRDA iPSCs after 2-3 months in culture. Our results indicate that in contrast to what was observed in DM1 cells, MSH2 does not play a role in maintaining DNA methylation at expanded FXS or FRDA alleles. In the course of doing this work, we also demonstrate that transcriptional activation of silenced *FMR1* alleles drives repeat contractions in FXS ESCs, some of which are MSH2-independent. These contractions may contribute to the repeat size mosaicism that is commonly seen in individuals who inherit *FMR1* alleles with large numbers of repeats. As such they may contribute to the variability in clinical presentation and disease penetrance that is observed.

## Methods

### Cell lines and culture conditions

Human FXS ESCs (WCMC37) were obtained from Nikica Zaninovic, Weill Cornell Medical College of Cornell University (New York, NY). The isolation and characterization of the 37F clone has been previously described (53). The FRDA iPSCs (GM23404) were obtained from Coriell Institute for Medical Research (Camden, NJ). Both FXS ESCs and FRDA iPSCs were cultured on tissue culture dishes coated with 5 µg/mL CellAdhere^TM^ Laminin-521 (STEMCELL Technologies, Cambridge, MA, 200-0117) in StemFlex^TM^ medium (Thermo Fisher Scientific, Waltham, MA, A3349401,) supplemented with 1X antibiotic-antimycotic (Thermo Fisher Scientific,15240096) at 37°C in 5% CO_2_. Medium was changed every other day. Cell passaging was performed twice a week using StemPro^TM^ Accutase^TM^ (Thermo Fisher Scientific, A11105-01) and cells were plated in growth medium supplemented with RevitaCell^TM^ supplement (Thermo Fisher Scientific, A2644501). Total RNA from H1 ESCs (WA01, WiCell Research Institute, Madison, WI) carrying typical *FMR1* alleles with 30 CGG repeats was used as a control for measuring *FMR1* mRNA levels and genomic DNA from GM06865 (Coriell Institute for Medical Research) lymphoblastoid cells that carry typical *FMR1* alleles, unmethylated with 30 CGG repeats was used as a control in the bisulfite sequencing for *FMR1* promoter methylation analysis. Genomic DNA from a human iPSC line carrying typical *FXN* alleles with 10 GAA repeats was used as a control in the bisulfite sequencing for methylation analysis of the *FXN* locus.

### Generation of MSH2 knockout cell lines using CRISPR/nCas9

Genome edits were made using nCas9 (D10A nickase mutant) to create a single-stranded break at two locations in the *MSH2* gene within exon 3. Repair by non-homologous end joining (NHEJ) resulted in random mutations, especially large >40 bp deletions, which led to complete knockout (KO) of MSH2. The nCas9 sgRNAs were previously described (39) and shown to have high editing efficiency in hESCs. To generate a fragment containing two complete sgRNA expression cassettes, oligonucleotide primers pX462-MSH2-dual-F and pX462-MSH2-dual-R (Table S1) were annealed to a U6-gRNA scaffold template containing gRNA-scaffold followed by U6 promoter as previously described (54). The PCR was done using Q5 Hot Start High-Fidelity DNA Polymerase (New England Biolabs, Ipswich, MA, M0493L) to generate a fragment that was cloned into a BbsI digested pSpCas9n(BB)-2A- Puro (PX462) V2.0, a gift from Feng Zhang (Addgene plasmid 62987; http://n2t.net/addgene:62987) (55) using the NEB HiFi DNA Assembly Cloning kit (New England Biolabs, E2621L). The ligated DNA was transformed into DH5alpha competent cells (New England Biolabs, C2987I) and plasmid DNA was isolated using the Monarch miniprep kit (New England Biolabs, T1010L). Vector assembly for pX462-MSH2-dual-gRNA was verified by Sanger sequencing (Psomagen, Rockville, MD), and transfection-grade plasmid DNA was prepared using the NucleoBond Xtra Midi Plus kit (Macherey-Nagel, Allentown, PA, 740422.5). Whole plasmid sequence is available in Additional file 8.

Human ESCs and iPSCs (1.7 x 10^5/ well) were plated in 12-well plates coated with Laminin-521 one day before transfection to be 50% confluent at time of transfection. Cells were transfected with pX462-MSH2-dual-gRNA and pCE-mp53DD, a gift from Shinya Yamanaka (Addgene plasmid # 41856; http://n2t.net/addgene:41856) (56) using Lipofectamine STEM (Thermo Fisher Scientific, STEM00008) according to the manufacturer’s guidelines. pCE-mp53DD was used to improve cell survival after CRISPR editing. Twenty-four hours after transfection, 1 µg/mL puromycin (Thermo Fisher Scientific, A11138-03) was added for 48 hours to select for transfected cells. After puromycin selection, cells were grown until well-separated colonies were visible and then passaged as single cells at low densities on 60 mm dishes coated with Laminin-521 and grown until colonies formed. Colonies were scraped and replated individually into 24 well plates.

Control cells were mock-transfected with only pCE-mp53DD plasmid. Single-cell colonies from the mock-transfected lines were picked to serve as wild type (WT) controls.

### Sequence validation of the MSH2 knockout

Exon 3 of the MSH2 gene was PCR amplified using Q5® Hot Start High-Fidelity DNA Polymerase with the following conditions: 98°C for 1 minute, (98°C for 15 seconds, 59°C for 15 seconds, 72°C for 40 seconds) x 35 cycles, and a final elongation step at 72°C for 2 minutes using 1 µM of Gb_HsMSH2_3F and Gb_HsMSH2_3R2 primers (Table S1). PCR products were either cleaned using ExoSAP-IT reagent (Thermo Fisher Scientific, 78201.1ML) or purified using NEB Monarch PCR cleanup kit. Cleaned PCR products were Sanger sequenced (Psomagen) and analyzed using the Synthego ICE software (Synthego Performance Analysis, ICE Analysis. 2019. v3.0. Synthego; [Accessed March 15, 2023]) to rule out the presence of a WT MSH2 exon 3 allele.

### Western blot analysis

To prepare total cell lysates, 1-2 x 10^6^ cells were washed once with cold PBS and resuspended in lysis buffer containing 10 mM Tris pH 7.5, 1mM EDTA pH 8.0, 1% Triton X- 100 (Sigma-Aldrich, St. Louis, MO, T8787-50ML), 1X protease inhibitor cocktail (Millipore Sigma, Burlington, MA, P8340-5ML), and 1X phosphatase inhibitor cocktail (Sigma-Aldrich, P5726-1ML). After 10 minute incubation on ice, lysate was sonicated in Bioruptor (Diagenode, Denville, NJ) on medium setting for 1 minute, 30s on/30s od and stored at - 80°C. Protein concentrations were measured using the Bradford Protein Assay Dye Reagent (Bio-Rad Laboratories, Hercules, CA, 5000006), and then 1X LDS loading buffer (Thermo Fisher Scientific, NP0007) and 1X Reducing Agent (Thermo Fisher Scientific, NP0009) were added to the lysate and heated for 5 min at 95°C. A total of 20 µg lysate was run on a NuPAGE^TM^ 4-12% Bis-Tris Gel (Thermo Fisher Scientific, NP0322BOX) in 1X MOPS SDS Running buffer (Thermo Fisher Scientific, NP0001) at 200V for 45 minutes. Proteins were transferred onto a nitrocellulose membrane using the Trans-Blot Turbo RTA transfer kit (Bio-Rad Laboratories, 1704270) and Trans-Blot Turbo Transfer system at 25V for 14 min.

Nitrocellulose membranes were blocked for 1 hour with 5% Amersham ECL Prime Blocking reagent (Millipore-Sigma, RPN418) in 1X TBST (Thermo Fisher Scientific, 28360). Blots were incubated overnight at 4°C in 1:10,000 diluted rabbit anti-human MSH2 antibody (Abcam, Waltham, MA, ab70270). The next day, blots were washed 3 times with 1X TBST for 5 min each and incubated for 1 hour at room temperature with 1:5000 diluted secondary anti- rabbit HRP conjugated antibody (Sigma Aldrich, GENA934-1ML). Blots were washed 3 times in TBST for 5 min each, and once in 1X TBS, and then detected with ECL prime western blot detection reagent (Millipore-Sigma, GERPN2232) for 5 minutes in dark. Bands were visualized using the Chemidoc Imaging System (Bio-Rad Laboratories). Blots were washed with 1X TBS and incubated with 1:1000 diluted anti-β-actin antibody (Abcam, ab8227) overnight at 4°C. The blot was then processed and imaged as before.

### Immunofluorescence

CRISPR edited FXS MSH2 KO-2 and FXS MSH2 WT-2 cells were plated in 24 well plates (2 x 10^5^ cells per well). The next day, cells were washed with PBS, fixed with 4% formaldehyde at 4°C for 45 minutes and washed three times with PBS. Cells were blocked in blocking buffer [PBS containing 10% normal goat serum (Thermo Fisher Scientific, PCN5000) and 0.3% Triton X-100 (Sigma-Aldrich, T8787-50ML)] for 1 hour. Fixed cells were then incubated with 1:1000 rabbit anti-human MSH2 antibody (Abcam, ab70270) diluted in PBS containing 1% BSA (Millipore Sigma, A7030-10G), 1% normal goat serum and 0.3% Triton X-100 at 4°C overnight. After three washes with 0.1% BSA in PBS, cells were incubated with 1:2000 secondary antibody (Thermo Fisher Scientific, A-11034) diluted in PBS containing 1% BSA, 1% normal goat serum and 0.3% Triton X-100 for 45 minutes. Nuclei were stained with DAPI (Thermo Fisher Scientific, 66248) diluted to 0.25 ug/mL in PBS with 0.1% BSA. Images were taken on BioTek Cytation 5 Cell Imaging Multimode Reader (Agilent, Santa Clara, CA) and processed with Fiji software (version 2.14.0/1.54f) (57).

### RNA methods

Total RNA was isolated from cells using TRIzol^TM^ reagent (Thermo Fisher Scientific) according to manufacturer’s instructions. RNA was quantified using DS-11 (DeNovix, Wilmington, DE). Three hundred nanograms of RNA was reverse transcribed to cDNA in 20 µL final volume using SuperScript^TM^ IV VILO^TM^ master mix (Thermo Fisher Scientific, 11756050) as per manufacturer’s instructions. Real-time PCR was performed in triplicate using 2 µL of the cDNA, FAM-labeled *FMR1* (Hs00924547_m1) and VIC-labeled *β- glucuronidase* (*GUSB*) endogenous control (4326320E) Taqman probe-primers (Thermo Fisher Scientific) and TaqMan Fast Advanced master mix (Thermo Fisher Scientific, 4444964) on QuantStudio 3 Real-Time PCR system (Thermo Fisher Scientific). For quantitation, the comparative threshold (ddCt) method was used.

### Analysis of CGG and GAA repeats via PCR

Genomic DNA was isolated by the salting out method (58). For *FMR1* CGG repeat PCR, 1 µg of genomic DNA from FXS cells was digested overnight at 37°C in rCutSmart Buffer (New England Biolabs, B6004S) with either only HindIII-HF (New England Biolabs, R3104S) or HindIII-HF and HpaII (New England Biolabs, R0171S). PCR was performed using 5 µL digested DNA in a 50 µL reaction containing 0.8 µM each of Not_FraxC and FAM- Not_FraxR4 primers (Table S1), 1X pH 9.0 Triton Buffer (50 mM Tris pH9, 1.5 mM MgCl2, 20 mM (NH_4_)_2_SO_4_, 0.2% Triton X-100), 2.8M betaine, 2.3% DMSO, 0.6 mM dNTPs and 0.5 units of Q5 Hot Start High-Fidelity DNA Polymerase using the following thermocycler conditions: 98°C for 3 minutes, (98°C for 30 seconds, 65°C for 30 seconds, 72°C for 3 minutes 30 seconds) x 33 cycles, and a final elongation step at 72°C for 10 minutes. For *FXN* GAA repeat PCR, ∼100 ng genomic DNA from FRDA cells was amplified using 0.5 µM each of GAA_104F and GAA_629R primers (Table S1) in a 20 µL reaction with 1X Q5 buffer, 0.2 mM dNTPs and 0.4 units Q5® Hot Start High-Fidelity DNA Polymerase with following thermocycler conditions: 98°C for 1 minute, (98°C for 15 seconds, 70°C for 15 seconds, 72°C for 3 minutes) x 35 cycles, and a final elongation step at 72°C for 10 minutes. For small pool *FMR1* CGG repeat PCR, a 200 µL master mix was assembled with 0.5 µM each of Not_FraxC and FAM-Not_FraxR4 primers (Table S1), 1X pH 9.0 Triton Buffer (50 mM Tris pH9, 1.5 mM MgCl2, 20 mM (NH_4_)_2_SO_4_, 0.2% Triton X-100), 2.5M betaine, 2% DMSO, 0.5 mM dNTPs, 4 units of Q5 Hot Start High-Fidelity DNA Polymerase and 20 ng of genomic DNA. The 100 µl master mix was then divided into 10 µl aliquots each containing ∼300 genomes and PCR performed using the following thermocycler conditions: 98°C for 3 minutes, (98°C for 30 seconds, 65°C for 30 seconds, 72°C for 3 minutes 30 seconds) x 35 cycles, and a final elongation step at 72°C for 10 minutes. Products were run on a 1% TAE agarose gel in 1X TAE buffer.

### Methylation-specific qPCR

To analyze DNA methylation at the *FMR1* promoter, methylation-specific qPCR was performed as previously described (53, 59). A total of 1 µg genomic DNA was diluted to 10 ng/µL in TE pH 8.0 and sonicated in Bioruptor for 5 min at medium setting, 30 seconds on/30 seconds od. To confirm that the fragment size was less that 1 kb, 15 µL of DNA was run on a 1% agarose gel in 1X TAE buffer. The sonicated DNA was divided into two tubes and was digested in one tube with HpaII restriction enzyme (New England Biolabs, R0171S) in rCutSmart Buffer (New England Biolabs, B6004S) overnight at 37°C. The second tube of undigested DNA was also incubated in parallel. The samples were incubated at 80°C for 20 minutes to inactivate HpaII. Real-time PCR was performed in triplicate using 2 µL each of undigested and digested DNAs using PowerUp^TM^ SYBR^TM^ Green Master mix (Thermo Fisher Scientific, A25777) using QuantStudio^TM^ 3 Real-Time PCR machine. The *FMR1* promoter was amplified using primer pair *FMR1* exon1 F and exon1 R, and *GAPDH* was amplified using primer pair *GAPDH* exon1 F and *GAPDH* intron1 R (Table S1). *GAPDH* was used as an unmethylated control region to confirm HpaII digestion. For quantitation, the comparative threshold (ddCt) method was used. Ct values of digested samples were compared to undigested samples to determine percent methylation. The extent of methylation determined by this assay sometimes exceeds 100% for fully methylated samples due to small differences in the Ct values resulting from pipetting error (59).

### Bisulfite Conversion and Sequencing

For the FXS samples, a total of 500 ng of genomic DNA was bisulfite-converted using the EZ DNA Methylation-Lightning Kit (Zymo Research, Irvine, CA, P212121) and eluted in 20 µL elution buffer. A hemi-nested PCR was performed to amplify the *FMR1* promoter region.

For the outer PCR, 4 µL bisulfite converted DNA was added to a PCR master mix to final concentrations of 0.25 µM each for primer (FMR1_Met_1F and Gb_Metbis_2645R) (Table S1), 0.167 mM dNTPs, 1X Platinum II PCR buffer, and 0.04 U/µL Platinum™ II Taq Hot-Start DNA Polymerase (Thermo Fisher Scientific, 14966001). The PCR was performed by following cycles: 94°C for 3 minutes, (94°C for 30 seconds, 50°C for 30 seconds, 72°C for 1 minute) x 20, and 72°C for 4 minutes. The product was diluted 1:10 for the inner PCR. The inner PCR uses same PCR conditions except with Gb_FMR1_Met_2F (Table S1) as the forward primer and 30 cycles.

For the FRDA samples, 2 µg of genomic DNA were digested for 3 hours with HindIII restriction enzyme (New England Biolabs, R0104S) and stored overnight at -20°C. Then, 200 ng of DNA was enzymatically bisulfite converted using the NEBNext® Enzymatic Methyl-seq Conversion Kit (New England Biolabs, E7125S) which produced 100 µL of converted DNA at 2 ng/µL concentration. The bisulfite modified DNA was stored overnight at 4°C. Then, 4 µL of DNA was used for PCR using 1.5 µM primers Gb_FXNMe-1212-F and Gb_ FXNMe_1930-R (Table S1, (22)), 0.2 mM dNTPs, 1X Platinum II PCR Buffer and 0.04 U/µL Platinum™ II Taq Hot-Start DNA Polymerase. The PCR was performed by following cycles: 95°C for 5 minutes, (95°C for 30 seconds, 59°C for 30 seconds, 72°C for 2 minutes) x 35, and 72°C for 10 minutes. The PCR products were cleaned using Zymo Select-a-Size DNA Clean and Concentrate (Zymo Research, D4084) and eluted in 20 µL NEB Elution Buffer (New England Biolabs, T1016L).

After determining the DNA concentration using the DS-11, bisulfite/enzymatic converted DNA samples were diluted to 7.5 ng/µL and analyzed by Plasmidsaurus Premium Sequencing (https://plasmidsaurus.com/home) which generated 500-3000 individual reads. These reads were aligned using NCBI BLAST software (60) to a bisulfite converted genomic DNA as template and analyzed using a custom Python3 script (Additional file 7) to determine the methylation status of the 37 individual CpGs (for *FMR1*) and 17 CpGs (for *FXN*) and average methylation of the bulk population. Sequences that did not align or completely span the PCR amplified region were discarded.

### Transient transfection of dCas9-TET1 for *FMR1* reactivation

Gibson assembly (61) was used to construct the plasmid pMLM3636-Puro-dCas9-TET1 using NEBuilder®HiFi DNA Assembly Master Mix (New England Biolabs, E2621L) and fragments derived from the following plasmids: PuroR from pSpCas9(BB)-2A-Puro (PX459) V2.0, a gift from Feng Zhang (Addgene plasmid # 62988; http://n2t.net/addgene:62988) (55), dCas9 from Inducible Caspex expression, a gift from Steven Carr & Samuel Myers (Addgene plasmid # 97421; http://n2t.net/addgene:97421) (62), and TET1CD from pPlatTET-gRNA2, a gift from Izuho Hatada (Addgene plasmid # 82559 ; http://n2t.net/addgene:82559) (63) and vector backbone from pMLM3636 (a gift from Keith Joung (Addgene plasmid # 43860 ; http://n2t.net/addgene:43860). Two additional c-MYC nuclear localization signal sequences (NLSs) were added using Gb_dCas9-cmyc-F (Table S1) to increase nuclear localization of dCas9-TET1. Various PCR fragments were purified with 0.5 volumes of Zymo Select-a-Size DNA Clean and Concentrator Magbead Kit (Zymo Research, # D4084) before assembly. The assembled DNA was transformed into DH5alpha competent cells (New England Biolabs, C2987I) and plasmid DNA was isolated using the Monarch miniprep kit (New England Biolabs, T1010L). Vector assembly was verified using Whole Plasmid Sequencing (Psomagen, Rockville, MD) and transfection-grade plasmid DNA was prepared using the NucleoBond Xtra Midi Plus kit (Macherey-Nagel, Allentown, PA, 740422.5).

To generate pMLM3636-Puro-dCas9-TET1-CGG, a GG(CGG)6 gRNA (gRNA-CGG_6_, Table S1) was cloned into pMLM3636-Puro-dCas9-TET1. To generate pMLM3636-Puro-dCas9-TET1- PRM with two different gRNAs in the promoter region (gRNA-PRM-1 and gRNA-PRM-2) previously described in (64), oligonucleotide primers PuroR-FMR1-gRNA-PRM_F and PuroR-FMR1-gRNA-PRM_R (Table S1) were annealed to a U6-gRNA scaffold template containing gRNA-scaffold followed by U6 promoter. The PCR was done using Q5 Hot Start High-Fidelity DNA Polymerase (New England Biolabs, Ipswich, MA, M0493L) to generate a fragment that was cloned into pMLM3636-Puro-dCas9-TET1. Whole plasmid sequences are provided in Additional Files 9, 10 and 11.

To transiently demethylate and reactivate the *FMR1* gene, human ESCs (8 x 10^5^/ well) were plated in 6-well plates one day before transfection to be 80% confluent at time of transfection. Cells were transfected with 1 µg of pMLM3636-Puro_dCas9-TET1-CGG6 and 200 ng of pCE-mp53DD using Lipofectamine STEM according to the manufacturer’s guidelines. Twenty-four hours after transfection, 1 µg/mL puromycin was added for 48 hours to select for transfected cells. Surviving cells were grown for 40 days under the cell culture conditions described above in one well of a 6-well plate and DNA and RNA were collected every 7 days for analysis.

### PCR to detect plasmid presence in transfected cells

The presence of transfected plasmid in cells at different days post transfection was tested by PCR. Either 200 ng of genomic DNA isolated from transfected cells, or 1 ng of plasmid (used as a positive control) was used in the PCR reaction along with 1X Q5 buffer, 0.04 mM dNTPs, 0.5 uM each of primers 3636-727F and 459-377R (Table S1) and 0.4 units Q5® Hot Start High-Fidelity DNA Polymerase. The following thermocycler conditions were used: 98°C for 1 minute, (98°C for 15 seconds, 60°C for 30 seconds, 72°C for 30 seconds) x 33 cycles, and a final elongation step at 72°C for 2 minutes. The PCR products were run on a 1% TAE agarose gel in 1X TAE buffer.

### Statistical methods

The statistical significance of the differences was calculated by student’s t-test using a cutoff of P <0.05 to determine significance.

## Results

### Loss of MSH2 does not aNect DNA methylation at the *FMR1* promoter in FXS ESCs

We used CRISPR/Cas9 genome editing with dual Cas9 nickases and 2 guide RNAs targeting human *MSH2* exon 3 to generate *MSH2*^-/-^ FXS ESCs (WCMC 37F) that carry a fully methylated *FMR1* gene with 400 CGG repeats (Figure 1A). A plasmid carrying a dominant negative version of p53, pCE-mp53DD, was co-transfected to increase cell survival. Exon 3 of *MSH2* encodes the DNA binding domain of the MSH2 protein, and the guide RNAs were chosen based on previous studies showing they generated efficient edits in hESCs (39). To generate control lines, FXS ESCs were mock transfected with pCE-mp53DD. To confirm the crispants, we sequenced a 631 base pair region containing all of MSH2 exon 3 and analyzed the mutations in both FXS MSH2 KO-1 and FXS MSH2 KO-2 using Synthego ICE software. We found complete loss of the parental allele in both cell lines caused by either large in-frame deletions or frameshift mutations (Figure S1A). The control cell lines, FXS MSH2 WT-1 and FXS MSH2 WT-2 showed no changes in the MSH2 exon 3 sequence (Figure S1A). The levels of MSH2 in edited lines were analyzed using a western blot (Figure S1B) and 2 independent clones that showed a complete loss of MSH2 were selected for further studies (FXS MSH2 KO-1 and FXS MSH2 KO-2). Similarly, two clones from mock transfected cells were selected as control cell lines (FXS MSH2 WT-1 and FXS MSH2 WT-2). FXS MSH2 KO-1 and FXS MSH2 KO-2 showed no MSH2 expression when compared to FXS MSH2 WT-1 and FXS MSH2 WT-2 (Figure 1B). Finally, we performed immunofluorescence for MSH2 and confirmed that MSH2 has strong nuclear expression in all FXS MSH2 WT-2 cells, while there was no detectable MSH2 expression in the crispant FXS MSH2 KO-2 cells (Figure 1C).

**Figure 1.**
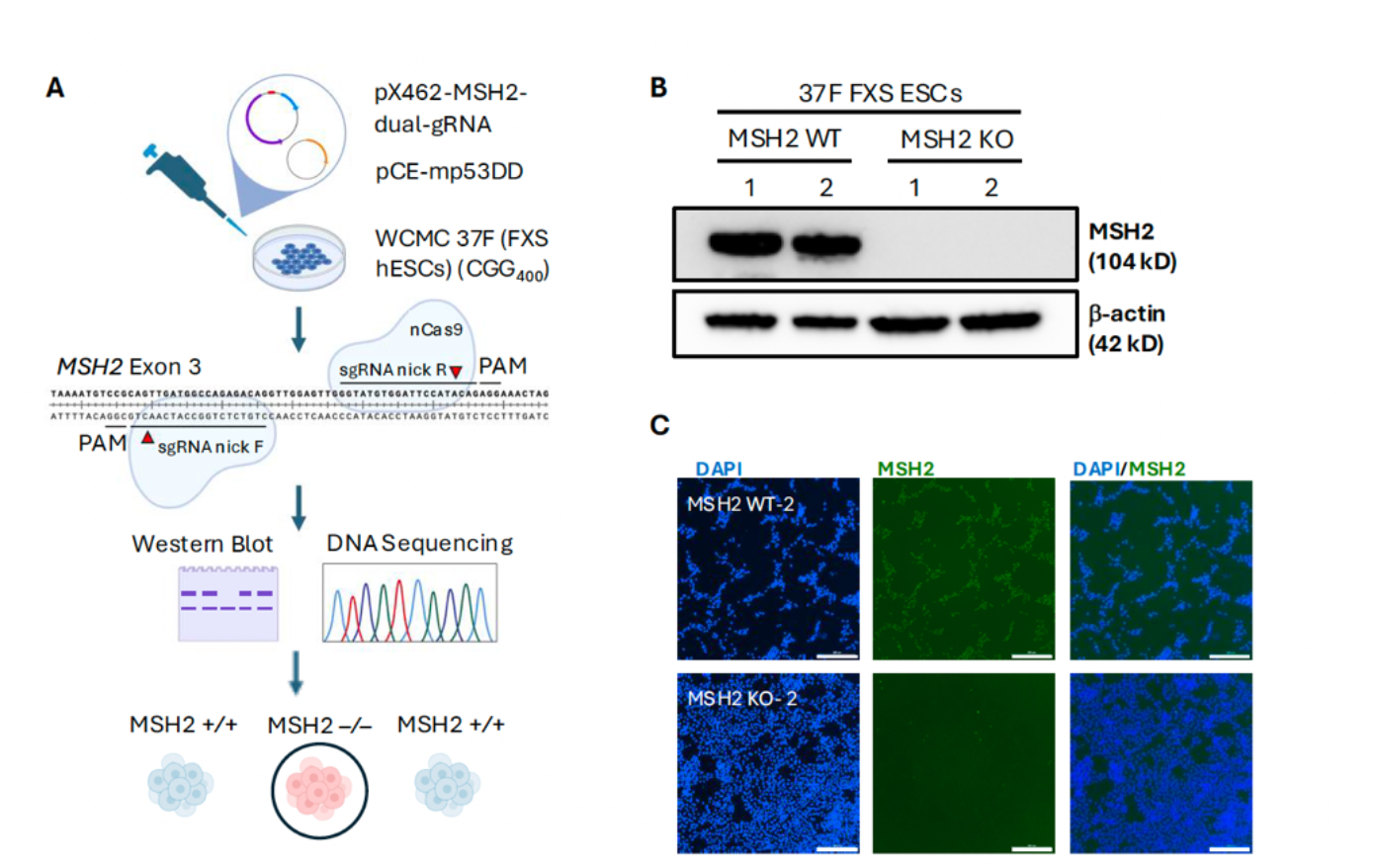
(A) A schematic showing the creation of MSH2 null fragile X syndrome human embryonic stem cell lines (FXS hESCs). WCMC 37F FXS ESCs were transfected with plasmids expressing nCas9 and dual guide RNAs targeting MSH2 exon 3. Edited clones were screened using western blot and DNA sequencing, and MSH2 null clones were selected for further study. **(B)** Western blot showing MSH2 expression in two cell lines each for FXS MSH2 WT and FXS MSH2 KO. β-actin used as loading control. Western blot of all clones assayed is shown in Figure S1. **(C)** Immunofluorescence imaging for MSH2 (green) shows a strong, nuclear signal in FXS MSH2 WT-2 and the absence of MSH2 in FXS MSH2 KO-2 cells. Scale bars represent 300 µm.

The two MSH2 WT and two MSH2 KO FXS ESC lines were grown for at least 100 days in culture and DNA methylation, *FMR1* mRNA levels and CGG repeat size were analyzed at indicated times for each line. We analyzed the methylation levels in the *FMR1* promoter for the two MSH2 WT and two MSH2 KO cell lines using a methylation-sensitive qPCR assay which provides a quantitative measure of bulk methylation. There was no significant difference in methylation between the MSH2 KO and MSH2 WT cell lines, indicating no change in the bulk methylation of the *FMR1* promoter more than 100 days post-MSH2 knockout (p = 0.75, Figure 2A). Next, we analyzed the *FMR1* mRNA levels as a proxy for any subtle decrease in DNA methylation. All lines expressed very low levels of *FMR1* mRNA, similar to those in the parental FXS ESCs from which these cell lines were derived, and no significant increase was seen in *FMR1* expression in either the MSH2 KO or MSH2 WT cell lines over 3 months in culture (Figure 2B). While human ESCs with a typical number of CGG repeats express *FMR1* mRNA at a level of 94% of *GUSB* expression (shown as a dotted line in Figure 2B), no FXS ESC line showed expression greater than 0.4% of *GUSB.* The MSH2 KO cell lines also did not show any obvious change in repeat size or DNA methylation when assessed by methylation-sensitive repeat PCR over 3 months in culture (Figure 2C). Thus, the loss of MSH2 did not have any significant effect on overall DNA methylation levels at the *FMR1* promoter in FXS ESCs.

**Figure 2.**
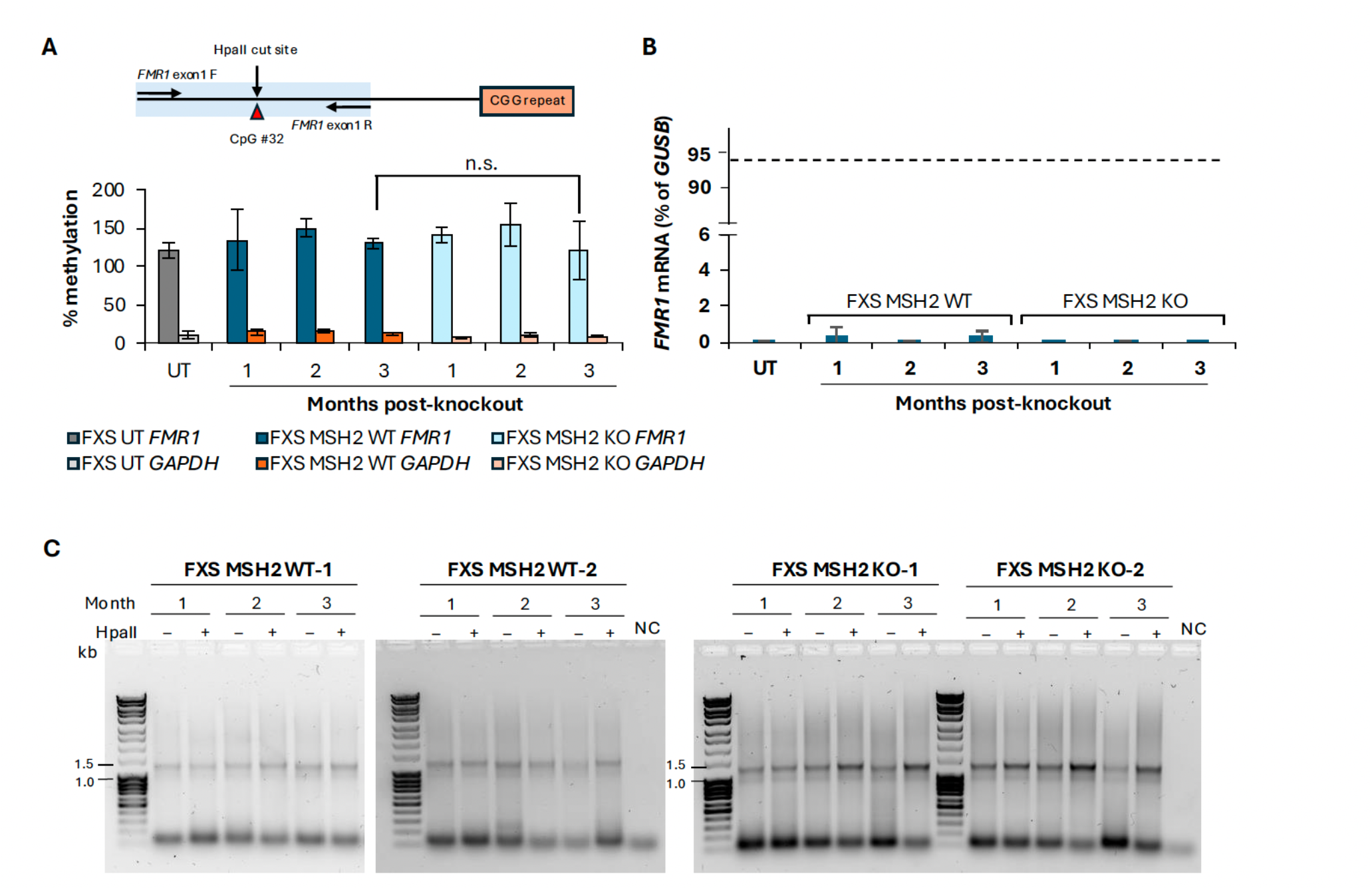
(A) Methylation qPCR in FXS MSH2 WT and FXS MSH2 KO ESCs. Above, schematic showing the location of qPCR primers and methylation-sensitive HpaII restriction enzyme site in the *FMR1* promoter region analyzed by the methylation qPCR assay. Below, % methylation data are shown as an average from two cell lines each for FXS MSH2 WT and FXS MSH2 KO and two replicates of untransfected (UT) FXS ESCs, and error bars represent standard deviation. *GAPDH* is used as an unmethylated region for HpaII digestion control. n.s., not significant. **(B)** *FMR1* mRNA levels in FXS MSH2 WT and FXS MSH2 KO ESCs are shown as a percentage of *GUSB* mRNA. Data shown are an average from two cell lines each for FXS MSH2 WT and FXS MSH2 KO and two replicates of untransfected (UT) cells, and error bars represent standard deviation. The dashed line shows the level of *FMR1* mRNA levels in H1 hESCs with typical number of CGG repeats at 94% of *GUSB*. **(C)** CGG-repeat PCR in two cell lines each for FXS MSH2 WT and FXS MSH2 KO is shown at indicated times in culture with and without predigestion of genomic DNA with methylation sensitive HpaII. NC, negative control. 1 month (approximately 40-50 days), 2 months (approximately 70-80 days) and 3 months (approximately 100-110 days).

In DM1 ESCs, it was observed that specific CpG residues in the *DMPK* locus became demethylated after 2 months in culture post-MSH2 knockdown, while other CpGs remained methylated (39). To test if MSH2 might have a similar effect on specific CpG residues in FXS ESCs, we performed bisulfite sequencing to look at the methylation status of 37 individual CpG sites in the *FMR1* promoter region from –394 to –93 bp upstream of the CGG repeat (Figure 3A). After bisulfite conversion of DNA, long-read sequencing was used to determine the methylation profiles of 500-3000 individual DNA molecules for each line. Minor variations in the methylation of CpGs across different alleles in the sample were observed in cells with a typical number of CGG repeats as well as in the parental FXS ESCs. As expected, in alleles with a typical number of CGG repeats, there was almost no methylation seen in the *FMR1* promoter region whereas in FX alleles the overall methylation of the region was greater than 90% (Figure 3B). There was a small increase in the average methylation of *FMR1* alleles in the MSH2 KO cell lines over time; however, this difference was not statistically significant (p = 0.61) (Figure 3C). Some *FMR1* alleles in FXS MSH2 WT lines were slightly less methylated to begin with, and there was a slight decrease in average methylation over time (Figure 3C). However, there was no significant difference in the overall methylation levels between the FXS MSH2 WT lines and the FXS MSH2 KO lines at 3 months post-knockout (p = 0.27, Figure 3C). This suggests that the slight changes in methylation seen over time in FXS MSH2 KO cell lines were due to random variation and not progressive demethylation. In support of this theory, the residues showing demethylation in each cell line did not show a consistent pattern (Figure 3D). There were no residues that were differentially methylated in FXS MSH2 KO ESCs when compared to FXS MSH2 WT ESCs, and regions showing demethylation at earlier time points were not demethylated at later time points, suggesting a lack of progressive demethylation over time. This variation indicates that FXS ESCs do not have 100% methylation at the *FMR1* promoter in every cell. Random CpG sites were unmethylated, and even highly unmethylated alleles were also present in the population (Figure 3D). In fact, most alleles were either highly methylated (>90%) or highly unmethylated (<10%). This is consistent with the suggestion that methylation and demethylation are progressive processes that, once initiated, lead to complete methylation or demethylation (65). However, unlike what was seen in DM1 ESCs, there was no significant loss of DNA methylation following MSH2 knockout in FXS ESCs, either across the promoter region or at specific CpG residues.

**Figure 3.**
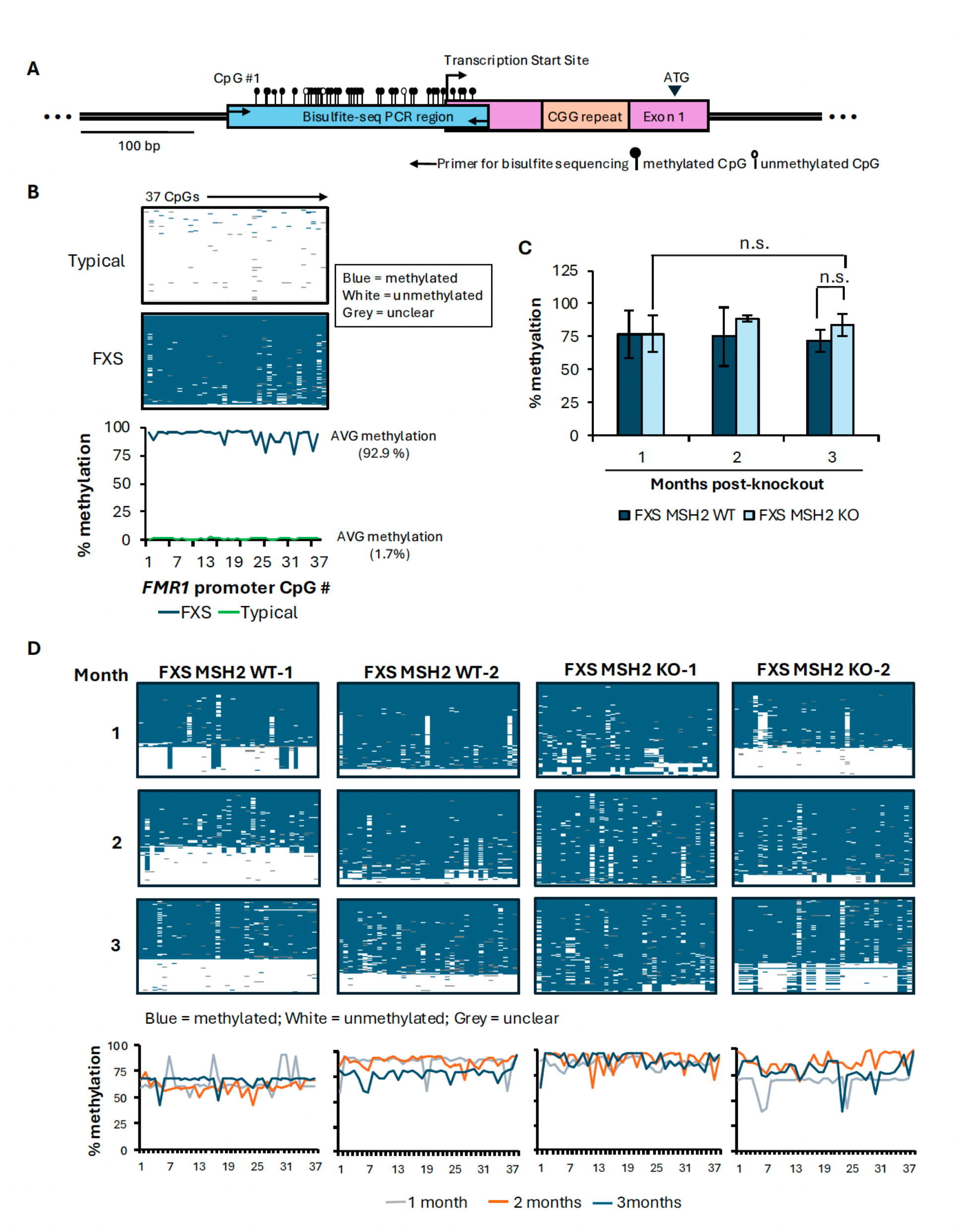
Bisulfite sequencing in FXS MSH2 WT and FXS MSH2 KO cell lines. (A) Schematic showing location of FMR1 promoter region assayed by bisulfite sequencing of 37 CpG residues. (B) Bisulfite sequencing data for individual CpG residues is shown from 100 random individual DNA reads arranged vertically from most to least methylated per DNA sample from a typical cell line with 30 CGG repeats (GM06865) used as an assay control, and untransfected WCMC 37F FXS ESCs. Summary panel below shows the average methylation at individual CpGs in the *FMR1* promoter region. (C) Average methylation in the *FMR1* promoter is shown for FXS MSH2 WT and FXS MSH2 KO cell lines at 1, 2 and 3 months. Bisulfite sequencing data for individual CpGs were averaged to compare overall methylation. Data shown is an average from 2 cell lines and error bars represent standard deviation. n.s., not significant. (D) Bisulfite sequencing data for individual CpG residues is shown from 100 random individual DNA reads arranged vertically from most to least methylated per DNA sample for 1, 2 and 3 months. Summary panels show the comparison of average methylation per CpG over time in each cell line.

### Loss of MSH2 does not aNect DNA methylation at the *frataxin* locus in FRDA iPSCs

In the *DMPK* gene, there are 25 CpG residues in the 318 bp region directly upstream of the CTG repeat, while the *FMR1* gene has 37 CpGs in the 349 bp region upstream of the CGG repeat (Figure S2A). However, the CGG repeats themselves contain 1 CpG per repeat, thus adding another 400 CpGs to the *FMR1* locus in the FXS ESCs used in the current study. To test whether the differences observed in the levels of DNA methylation upon loss of MSH2 between *DMPK* alleles in DM1 ESCs and *FMR1* alleles in FXS ESCs could be attributed to the differences in the overall CpG density, we decided to study the role of MSH2 in another repeat expansion disorder, FRDA, because the differentially methylated region upstream of expanded GAA repeat in *FXN* alleles is less CpG-rich (Figure S2) and the GAA repeat itself cannot be methylated. We generated two *MSH2* knockout clones (FRDA MSH2 KO-1 and FRDA MSH2 KO-2) and two mock-transfected control clones (FRDA MSH2 WT-1 and FRDA MSH2 WT-2) in an FRDA iPSC line (GM23404), which has ∼700-800 GAA repeats in the *FXN* gene. We used the same dual nickase strategy and guide RNAs used to generate the *MSH2* knockout in FXS ESCs (Figure 1A). We sequenced *MSH2* exon 3 PCR products from the FRDA MSH2 KO clones, analyzing them using Synthego ICE software (Figure S3A). We found that both FRDA MSH2 KO cell lines had complete loss of the parental *MSH2* alleles due to large insertions or deletions. We further confirmed the loss of MSH2 via western blot (Figure 4A and Figure S3B).

**Figure 4.**
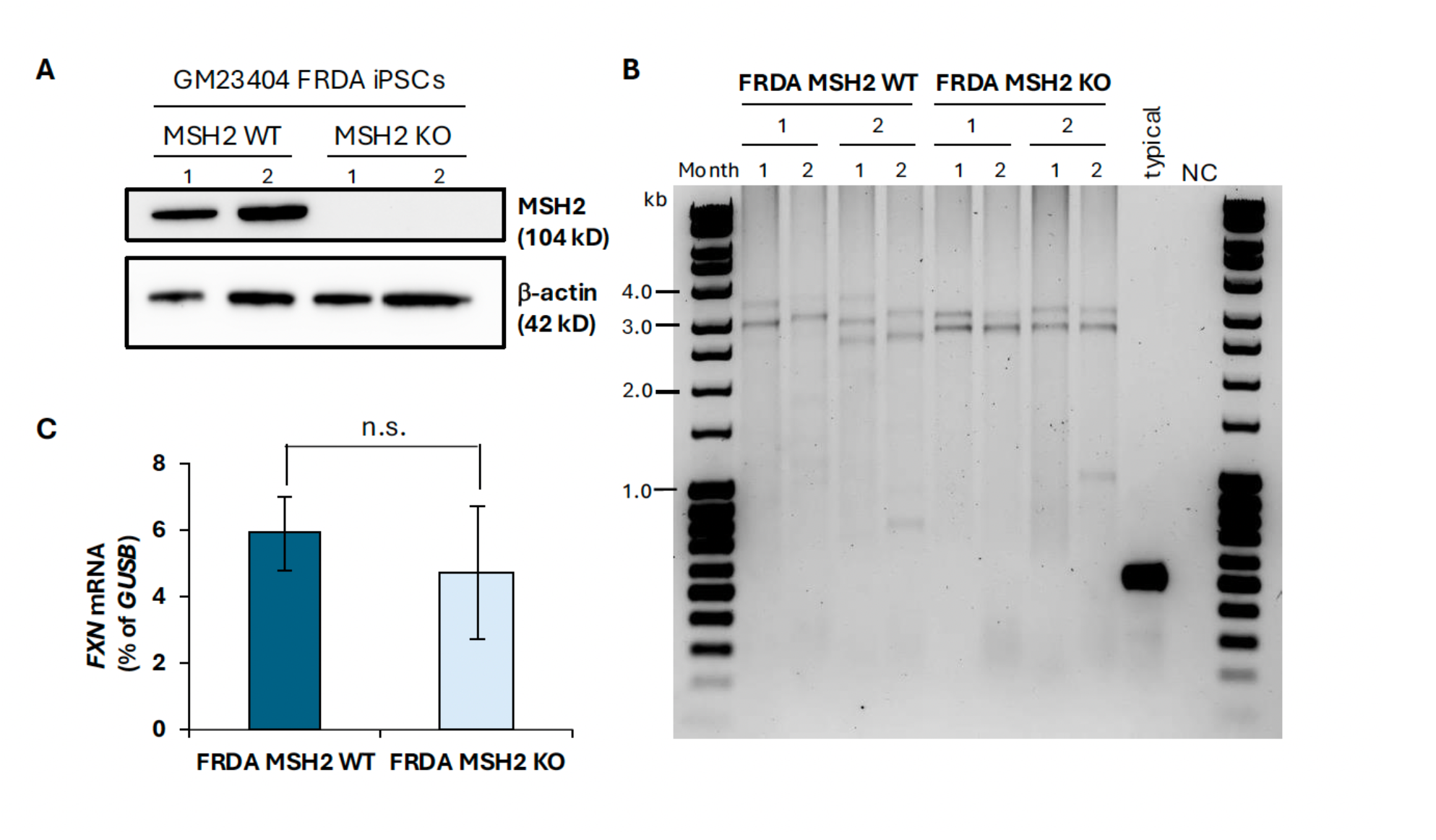
Validation of MSH2 CRISPR knockout in FRDA iPSCs. **(A)** Western blot showing MSH2 expression in FRDA MSH2 WT and its loss in FRDA MSH2 KO cell lines. **(B)** PCR for GAA repeat size for FRDA MSH2 WT-1 and WT-2, and FRDA MSH2 KO-1 and KO-2 cells at 1 and 2 months in culture. Genomic DNA from WCMC 37F cells which carry *FXN* alleles with ∼10 GAA repeats was used as a positive control (typical) and water as negative control (NC). **(C)** Average *FXN* mRNA levels from two FRDA MSH2 WT and two FRDA MSH2 KO lines measured at 2 months post knockout. Data shown are an average from two independent experiments and the error bars represent standard deviation. n.s., not significant.

The MSH2 WT and MSH2 KO FRDA iPSC lines were grown for at least two months under the same conditions as the FXS ESCs. To further characterize the lines, we performed GAA repeat PCR. Each cell line had slightly different GAA repeat sizes, but all were within the range of 700-850 repeats, reflecting the variability in repeat size in the parental population (Figure 4B). We observed that over two months in culture, the FRDA MSH2 WT-1 and FRDA MSH2 WT-2 lines showed visible expansion: for each cell line, both alleles increased in size between one month and two months post mock-transfection. In FRDA MSH2 KO-1 and FRDA MSH2 KO-2, there was no increase in allele size over two months post-*MSH2* knockout. This is consistent with previous studies that showed that MSH2 is required for repeat expansion in FRDA iPSCs (35). In both FRDA MSH2 WT and FRDA MSH2 KO cell lines, the GAA repeat PCR showed bands indicating stochastic contractions within the population, which did not always persist. This had been previously observed in FRDA iPSC lines, but interestingly, while loss of MSH2 was previously shown to decrease contractions (35), our data show that loss of MSH2 did not completely eliminate them (Figure 4B). We also measured *FXN* mRNA levels over time in the four FRDA iPSC lines. However, there was no consistent trend over time, and no significant difference in *FXN* mRNA levels was seen between FRDA MSH2 WT and FRDA MSH2 KO cell lines at two months post-transfection (p = 0.55, Figure 4C).

Next, we used bisulfite sequencing to assay the 718 bp region from -816 to -98 bp upstream of the GAA repeat in intron 1 of *FXN*, which contains 17 CpG sites (Figure 5A), a CpG density about 4 times lower than that of *FMR1* (Figure S2A). The parental GM23404 iPSCs showed high methylation at all 17 CpGs, with a higher average methylation at CpGs closer to the repeat. In comparison, an iPSC line from an individual with GAA repeats in the typical range showed differentially demethylated residues for CpGs 1-5, 8-9, 13 and 15 (Figure 5B), which includes 3 CpGs (5, 8 and 15) previously found to be unmethylated in typical alleles compared to FRDA alleles in lymphoblastoid cells (22). We observed that these additional residues are also differentially methylated in FRDA iPSCs when compared to typical iPSCs (Figure 5B). A similar pattern of differential methylation was previously seen in peripheral blood mononuclear cells (23). In the FRDA MSH2 WT lines, this region was highly methylated at each individual CpG (>90%) and methylation did not change significantly over time (Figure 5C). Similarly, in FRDA MSH2 KO lines, methylation in this region remained at >90% after two months post-MSH2 knockout and there was no significant difference in the DNA methylation levels between the FRDA MSH2 WT and FRDA MSH2 KO lines (p = 0.72, Figure 5C). No specific CpG residues that lost methylation over time in FRDA MSH2 KO iPSC lines were observed (Figure 5D). We did note that methylation was denser in the region closer to the GAA repeats, whereas CpG residues further from the repeats (CpGs #1-5) were slightly less methylated on average in both FRDA MSH2 WT and FRDA MSH2 KO cell lines (Figure 5B and Figure 5D).

**Figure 5.**
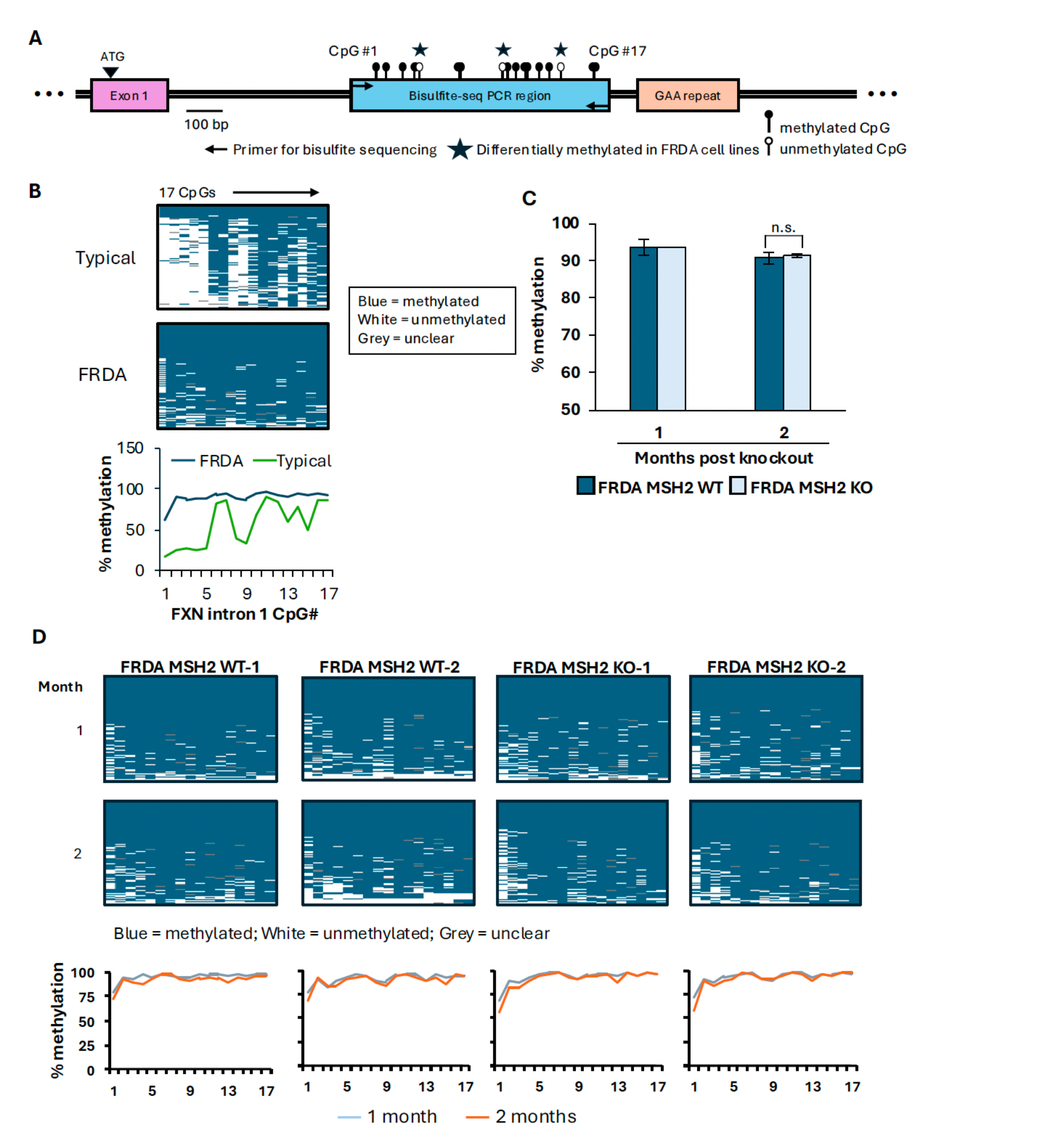
Bisulfite sequencing results for FRDA MSH2 WT and FRDA MSH2 KO cells. **(A)** Diagram indicating locations of the 17 CpG residues studied by bisulfite sequencing in the region 5’ of the GAA repeat in *FXN* intron 1. Residues marked with stars were found to be differentially methylated in FRDA cells compared to typical cells. **(B)** Bisulfite sequencing results show methylation at 17 individual CpGs in parental GM23404 human FRDA iPSCs and a human iPSC line carrying *FXN* alleles with typical GAA repeats. Data from individual CpG residues is shown from 100 random individual DNA reads arranged vertically from most to least methylated per genomic DNA sample from a human iPSC line with typical number of GAA repeats used as an assay control, and untransfected GM23404 FRDA iPSCs. Summary panel below shows average methylation at individual CpGs in the *FXN* promoter region. **(C)** Bisulfite sequencing at individual CpGs was averaged to compare overall methylation in FRDA MSH2 WT and FRDA MSH2 KO conditions over 2 months post MSH2 CRISPR knockout. Data shown is from an average of 2 cell lines and error bars represent standard deviation. n.s., not significant. **(D)** 17 CpG residues from 100 individual DNA sequences from bisulfite sequencing arranged vertically from most to least methylated per DNA sample. Summary panels below show the comparison of average methylation per CpG over time in each cell line.

### MSH2’s role in *de novo* methylation of *FMR1* alleles is unclear

While MSH2 transgene re-expression in DM1 MSH2 knockdown cell lines rescued CTG repeat instability, it failed to increase DNA methylation even when the repeat number exceeded the original, indicating that MSH2 was not involved in *de novo* methylation at the *DMPK* locus (39). Our observations for the *FMR1* locus in MSH2 KO FXS ESCs and the *FXN* locus in MSH2 KO FRDA iPSCs clearly indicate that MSH2 is not required for the maintenance of DNA methylation at these regions. However, since we did not see a decrease in DNA methylation with MSH2 KO, we could not use the MSH2 transgenic re-expression strategy to study its role in *de novo* methylation in either FRDA iPSCs or FXS ESCs.

Previous work has shown that the *FMR1* gene can be transiently demethylated and reactivated in FXS lymphoblastoid and fibroblast cells by treatment with 5- azadeoxycytidine (AZA), and the gene is re-silenced in approximately three weeks after removal of AZA (31, 66). We have previously used this strategy to study the role of PRC2 complex in *FMR1* gene silencing (49). However, given the toxicity of AZA treatment, we could not use this strategy to evaluate the role of MSH2 in *de novo* DNA methylation of *FMR1* alleles in FXS ESCs. We therefore used transient expression of ten-eleven translocation 1 (TET1) to reactivate the gene. TET1 catalyzes the first step in the demethylation of 5mC in CpGs (67, 68) and has been previously used to reactivate the *FMR1* gene in FXS cells (69, 70, 71). In one of these studies (69), massive contractions of CGG repeats were observed 27 days after lentivirus-mediated expression and targeting of TET1 to CGG repeats in FXS ESCs. However, these contractions were reported to be dependent on MSH2 (69). We reasoned that given the requirement of MSH2 for repeat contractions in FXS ESCs, transient plasmid-derived expression of TET1 in MSH2 KO FXS ESCs might allow us to reactivate the gene without generating contractions. Therefore, we transiently transfected FXS MSH2 WT-2 and FXS MSH2 KO-2 cell lines with plasmids expressing dCas9-TET1 and a guide RNA targeting the CGG repeats (gRNA-CGG). We measured *FMR1* mRNA levels by RT-qPCR and DNA methylation by methylation-specific qPCR over 40 days in culture (Figure 6). However, because of the reduced survival of transfected cells after puromycin selection (likely due to a low efficiency of plasmid transfection), the earliest time point at which we could collect the cells was at day 7 for FXS MSH2 WT cells and day 18 for FXS MSH2 KO cells. We observed the expected increase in *FMR1* mRNA (Figure 6A) and decrease in DNA methylation (Figure 6B) in both FXS MSH2 WT and FXS MSH2 KO cells; however, the gene remained unmethylated and active for more than 37 days in culture expressing *FMR1* mRNA at levels similar to or higher than that observed in H1 ESCs carrying *FMR1* alleles with the typical number of CGG repeats. This was not due to the presence of residual dCas9-TET1 plasmid (Figure S4). Rather, repeat size analysis showed that in both MSH2 WT-2 and MSH2 KO-2 cell lines, the band corresponding to the 400-CGG repeat was almost completely gone, replaced by a smear of unmethylated alleles with a wide range of repeat contractions (Figure 6C). Digestion of the template prior to PCR using a methylation-sensitive enzyme HpaII with recognition sites within the PCR amplicon revealed the presence of fully methylated large alleles in some samples. However, these were such a small proportion of the total allele population that they were not amplified unless the unmethylated alleles were digested and eliminated. The absence of a full-length PCR product was not due to the poor quality of genomic DNA as the small pool PCR on the same DNA sample showed contracted alleles of similar sizes in both MSH2 WT and MSH2 KO cells (Figure 6D).

**Figure 6.**
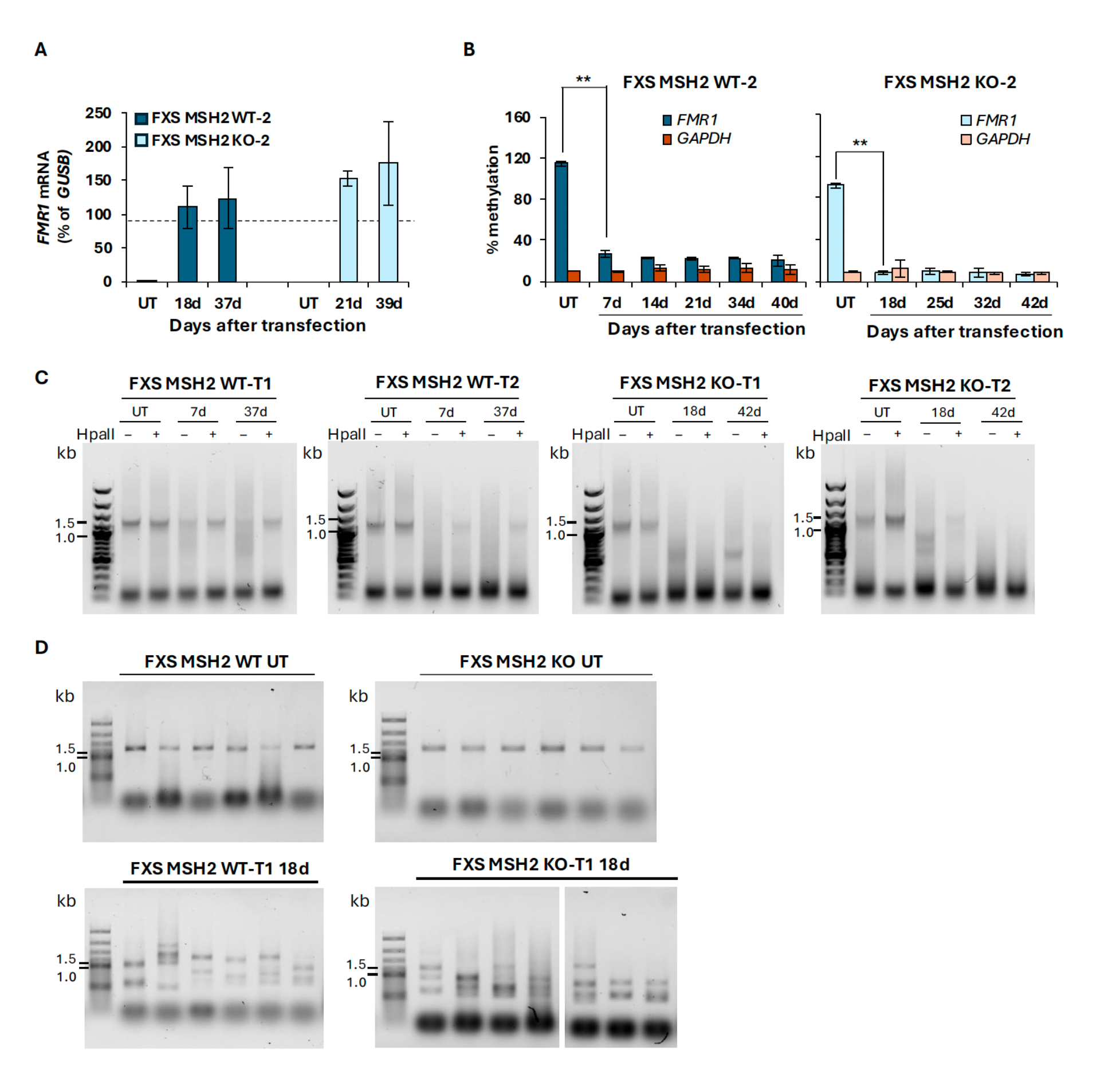
Transient transfection of FXS MSH2 WT-2 and FXS MSH2 KO-2 cell lines with plasmid pMLM3636-Puro-dCas9-TET1-CGG. **(A)** *FMR1* mRNA levels are shown for FXS MSH2 WT-2 and FXS MSH2 KO-2 cells untransfected (UT) and at different days post transfection with plasmid encoding dCas9-TET1 and gRNA targeting the CGG repeats. Data shown are an average of two independent experiments and error bars represent standard deviation. The dashed line represents *FMR1* mRNA levels in H1 ESCs with typical number of CGG repeats at 94% of *GUSB*. **(B)** Methylation-sensitive qPCR results for the *FMR1* promoter after dCas9-TET1-CGG_6_CG transfection in FXS MSH2 WT-2 and FXS MSH2 KO-2 cells. *GAPDH* is used as an unmethylated region for HpaII digestion control. Data shown are an average of two independent experiments and error bars represent standard deviation. ** p = 0.001. **(C)** CGG repeat PCR on DNA from two different transfections (T1 and T2) in FXS MSH2 WT-2 and FXS MSH2 KO-2 cells with plasmid dCas9-TET1-CGG. HpaII digested DNA lanes show amplification of alleles that are fully methylated while undigested samples amplify both methylated and unmethylated alleles. The *FMR1* gene in FXS ESCs carries 400 repeats resulting in a PCR product of about 1.4 kb. FXS MSH2 WT-2 and FXS MSH2 KO-2 cells show CGG repeat contractions indicated by discrete bands and smears below 1 kb. FXS MSH2 KO cells show almost complete loss of methylated full- length alleles in both transfection replicates. **(D)** Small pool PCR on the same DNA sample used for PCRs in panel C shows full length PCR product in untransfected (UT) cells and contracted alleles in both MSH2 WT and MSH2 KO cells transfected with plasmid dCas9- TET1-CGG.

Transfection with dCas9-TET1 either alone or with a dual guide targeting the *FMR1* promoter (gRNA-PRM-1 and gRNA-PRM-2) did not lead to any significant gene reactivation in either FXS MSH2 WT-2 or FXS MSH2 KO-2 cell lines (p = 0.1 for both, Figure 7A), likely due to insufficient TET1-targeting as evidenced by a lack of demethylation of the *FMR1* promoter (Figure 7B). There were no significant changes seen in the repeat size upon transfection with dCas9-TET1 either alone or with gRNAs targeting the *FMR1* promoter in both bulk and small pool PCR (Figures 7C, 7D and S5).

**Figure 7.**
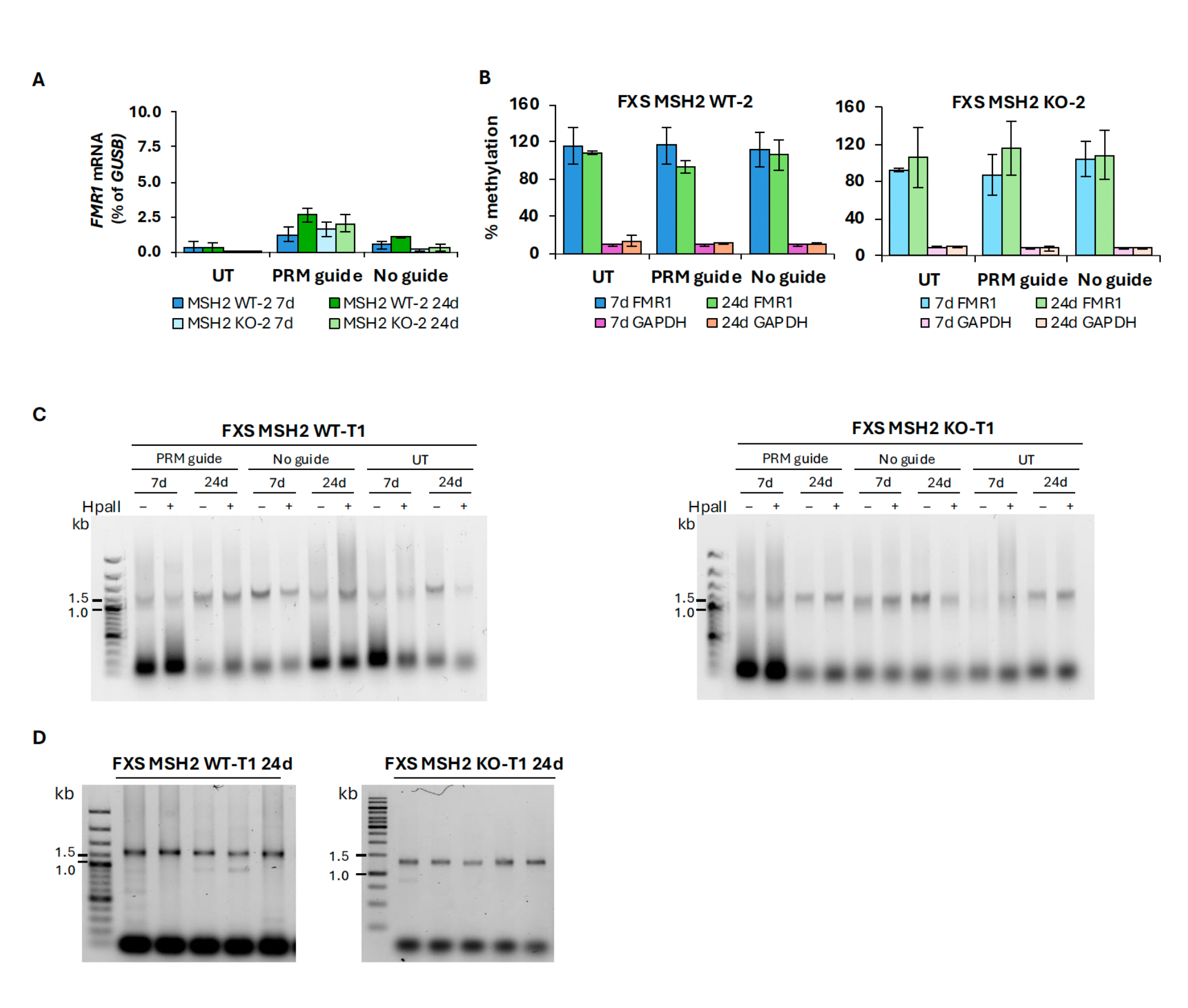
Transient transfection of FXS MSH2 WT-2 and FXS MSH2 KO-2 cell lines with plasmid pMLM3636-Puro-dCas9-TET1-PRM and pMLM3636-Puro-dCas9-TET1. **(A)** *FMR1* mRNA levels in FXS MSH2 WT-2 and FXS MSH2 KO-2 cells are shown as a percentage of *GUSB* mRNA in untransfected (UT) cells and at day 7 (7d) and day 24 (24d) after transient transfections with plasmid dCas9-TET1-PRM with 2 gRNAs directed to the *FMR1* promoter or with plasmid dCas9-TET1 with no gRNA. Data shown are an average of two independent experiments and error bars represent standard deviation. **(B)** Methylation-sensitive qPCR for the *FMR1* promoter at day 7 (7d) and day 24 (24d) after transient transfection with plasmids expressing dCas9-TET1 and gRNAs targeted to the *FMR1* promoter or dCas9-TET1 with no guide. *GAPDH* is used as an unmethylated region for HpaII digestion control. Data shown are an average of two independent experiments and error bars represent standard deviation. **(C)** CGG repeat PCR results for DNA from untransfected (UT) cells and replicate 1 (T1) at 7d and 24d for cells transfected with plasmid dCas9-TET1-PRM and plasmid dCas9-TET1 without any gRNAs. HpaII-digested DNA lanes show amplification of alleles that are fully methylated while undigested samples amplify both methylated and unmethylated alleles. The *FMR1* gene in FXS ESCs carries 400 repeats resulting in a PCR product of about 1.4 kb. FXS MSH2 WT and FXS MSH2 KO cell lines transfected with dCas9- TET1-PRM and dCas9-TET1 with no guide show CGG repeat sizes similar to untransfected (UT) cells at day 7 (7d) and day 24 (24d). The results of CGG repeat PCR for a replicate transfection (T2) are shown in Figure S5. **(D)** Small pool PCR on the same DNA sample used for PCRs in panel C shows full length PCR product in both MSH2 WT and MSH2 KO cells transfected with plasmid dCas9-TET1-PRM.

## Discussion

To test whether the role of MSH2 in the maintenance of DNA methylation seen at the *DMPK* locus in DM1 ESCs was conserved in two other REDs, FXS and FRDA, we made null mutations in *MSH2* in FXS ESCs and FRDA iPSCs and examined the effect on DNA methylation of *FMR1* and *FXN* alleles respectively. Our results indicate that MSH2 does not play a role in maintaining DNA methylation at either the *FMR1* locus in FXS ESCs or the *FXN* locus in FRDA iPSCs. The observed difference in MSH2-dependence is not due to the difference in the CpG-density because although the *FMR1* locus has a higher CpG-density than the *DMPK* locus, the *FXN* locus does not (Figure S2). Thus, more than one mechanism is likely involved in the maintenance of DNA methylation near expanded repeats with different pathways being utilized at different loci. It is worth noting here that MSH2 was not required to maintain the methylation of differentially methylated CpGs in the region downstream of the expanded CTG repeats in DM1 ESCs (39).

We could not use the MSH2 transgene re-expression strategy to evaluate the role of MSH2 in *de novo* methylation in FXS and FRDA cells as we did not observe a decrease in DNA methylation upon MSH2 knockout. Furthermore, we were also unable to test the effect of MSH2 on *de nov*o methylation in FXS cells after TET1 mediated demethylation of the *FMR1* promoter because, in contrast to a previous report (69), reactivation of FXS alleles resulted in a high frequency of contractions even in cells lacking MSH2 (Figure 6C). The contractions in both WT and *MSH2* null lines gave rise to *FMR1* alleles with repeat sizes that were below the threshold for *de novo* methylation (14, 53, 72). Thus, transcription drives at least two types of CGG repeat contractions, with a significant number of these contractions being MSH2-independent.

We and others have shown that actively transcribed *FMR1* alleles form R-loops that become more stable with increasing repeat number (49, 50, 73). R-loops are a frequent source of DNA damage, in part because the single-stranded DNA associated with the R-loop is prone to a variety of types of DNA damage even in the absence of replication (74, 75, 76). Furthermore, stable R-loops are also a frequent source of transcription-replication collisions. As such the R-loops formed on transcribed FX alleles may represent source of significant transcriptional stress that results in the contractions seen on reactivation of silenced FX alleles. A transcription-coupled repair (TCR) process has been implicated in CAG-repeat contractions in human cells (77). However, this process is MSH2-dependent.

In yeast, CAG-repeats have been shown to undergo transcription-induced contractions via an MSH2-independent process (78). Whether a similar mechanism is responsible for the MSH2-independent CGG-contractions we observe remains to be seen. A better understanding of the mechanisms underlying transcription induced repeat contractions may allow these processes to be harnessed to safely drive alleles from above a pathological repeat size threshold into or closer to the typical repeat size. The fact that transcription drives these contractions so effectively lends support to the idea that transcriptional silencing may be a cellular response to limit the risk to genome integrity.

Interestingly, very little demethylation and reactivation was observed when TET1 was overexpressed alone or when it was targeted to the *FMR1* promoter region (Figures 7A and 7B), and no repeat instability was seen in either FXS MSH2 WT-2 or FXS MSH2 KO-2 cells (Figure 7C and Figure S5) suggesting that TET1 activity at the CGG repeats is necessary for demethylation of the entire *FMR1* promoter region, gene reactivation and CGG repeat contractions.

## Conclusions

We have demonstrated that loss of MSH2 does not affect DNA methylation at the *FMR1* promoter in FXS ESCs or in intron 1 of the *FXN* gene in FRDA iPSCs. While we could not evaluate the role of MSH2 in establishing DNA methylation at the *FMR1* locus in FXS ESCs, our results clearly show that MSH2 is not required for the maintenance of DNA methylation induced by repeat expansion in either FXS or FRDA, and that this difference from what is seen in DM1 cannot be attributed to the difference in the overall GC content or CpG density of the regions examined. We suggest that any difference in the requirement of factors in the establishment or maintenance of repeat induced epigenetic changes may be due to the differences in the sequence or the structures formed by the expanded repeats. More work is needed to understand what these differences might be. Our work also revealed the existence of an MSH2-independent pathway for transcription-induced contractions. These contractions may contribute to the repeat size mosaicism seen in individuals with transcriptionally active unmethylated FX alleles that can influence the clinical presentation of the disease (79, 80, 81).

## List of abbreviations

RED: repeat expansion disorder
DM1: myotonic dystrophy type 1
FXS: fragile X syndrome
ESC: embryonic stem cell
FRDA: Friedreich’s ataxia
UTR: untranslated region
ASD: autism spectrum disorder
MMR: mismatch repair
hESC: human embryonic stem cell
iPSC: induced pluripotent stem cell

## Declarations

### Ethics approval and consent to participate

Not applicable.

## Consent for publication

All authors read and approved the manuscript.

## Availability of data and materials

All data generated in this study are included in the manuscript and additional files. Materials used in the current study are available from the corresponding author on reasonable request.

## Authors’ Contributions

Study conception and design: DK and KU; Plasmids and cell line constructions: JGB, KR and BH; Data collection: JGB and KR; Data analysis and interpretation: JGB, KR, BH, DK and KU; Manuscript writing, original draft: JGB, DK; Manuscript editing and revision: all authors; Supervision: DK; Funding and resources: KU

## Acknowledgements

Figures were created with Biorender. We would like to thank the other members of the Usdin lab, especially Carson Miller and Xiaonan Zhao for project advice and mentorship.

## Additional Files

Additional File 1: **Table S1** listing primers and gRNA sequences.

Additional file 2: **Figure S1. (A)** Sanger Sequencing results for MSH2 exon 3 PCR on FXS MSH2 WT and FXS MSH2 KO cell lines aligned to MSH2 exon 3. Diagram shows location of nickase CRISPR guide RNAs and single strand break locations. FXS MSH2 WT samples show 100% match to reference sequence and both FXS MSH2 KO samples display sequence breakdown within the two CRISPR nick sites indicative of CRISPR editing and mixed allele population. Sequence analysis using Synthego ICE software shows complete loss of *MSH2* exon 3 allele in both cell lines, with indel regions that match the region of sequence breakdown. **(B)** Western blot for MSH2 with β-actin loading control on single-cell clones of WCMC 37F FXS ESCs edited with dual CRISPR nickase targeting *MSH2* exon 3. All clones were analyzed here, and complete knockouts A6 and D3 (marked with asterisks) were selected for further study and renamed FXS MSH2 KO-1 and FXS MSH2 KO-2.

Additional file 3: **Figure S2. (A)** Diagram showing CpG residues that are methylated in regions upstream of the disease-causing repeats in Fragile X syndrome (*FMR1),* Friedreich’s Ataxia (*FXN),* and Myotonic Dystrophy Type I (*DMPK).* CpGs that are differentially methylated in FRDA vs non-FRDA lymphoblastoid cell lines (22) are noted with an asterisk and numbered. **(B)** Overall GC content of the region 1 kb upstream of the repeats in each disorder. The height of the curve and the color of the area under the curve are based on the average GC content (%) from low (blue) to high (red).

Additional file 4: **Figure S3. (A)** Sanger Sequencing results for MSH2 exon 3 PCR on FRDA MSH2 KO cell lines aligned to MSH2 exon 3. Diagram shows location of nickase CRISPR guide RNAs and single strand break locations. Both FRDA MSH2 KO samples show evidence of CRISPR editing: FRDA MSH2 KO-1 has an 80 bp insertion of a region of *MSH2* intron 3 inserted in between the CRISPR cut sites and FRDA MSH2 KO-2 has a 41 bp deletion with the 3’ cut site matching the sgRNA nick R cut site. Sequence analysis using Synthego ICE software shows complete loss of *MSH2* exon 3 allele in both cell lines, with a deletion matching the Sanger sequencing alignment in FRDA MSH2 KO-2. In FRDA MSH2 KO-1, ICE did not identify the insertion, but it did confirm complete loss of the typical *MSH2* allele. **(B)** Western blot for MSH2 with β-actin loading control on single-cell clones of GM23404 FRDA human iPSCs edited with dual CRISPR nickase targeting *MSH2* exon 3. All clones isolated are analyzed here and 4 clones were found to be complete knockouts: D1.1, C6.2, C4.1, D6.1. We selected two clones C6.2 and D1.1 for continued study (marked with asterisk in the Figure) and renamed them FRDA MSH2 KO-1 and FRDA MSH2 KO-2.

Additional file 5: **Figure S4**. PCR showing that low levels of TET1 plasmids are present at day 7 after transfection and completely lost by 2 weeks post transfection.

Additional file 6: **Figure S5**. CGG repeat PCR results for DNA from replicate transfection (T2) at 7d and 24d for FXS MSH2 WT-2 and FXS MSH2 KO-2 cells transfected with plasmid dCas9-TET1-PRM and plasmid dCas9-TET1 without any gRNAs. HpaII-digested DNA lanes show amplification of alleles that are fully methylated while undigested samples amplify both methylated and unmethylated alleles. The *FMR1* gene in FXS ESCs carries 400 repeats resulting in a PCR product of about 1.4 kb. FXS MSH2 WT and FXS MSH2 KO cell lines transfected with dCas9-TET1-PRM and dCas9-TET1 with no guide show CGG repeat sizes similar to untransfected (UT) cells at day 7 (7d) and day 24 (24d). The results of CGG repeat PCR for transfection T1 are shown in Figure 7C.

Additional file 7: Methylation analysis script .py

Additional file 8: Whole plasmid sequence for pX462-MSH2-dual-gRNA.dna

Additional file 9: Whole plasmid sequence for pMLM3636-Puro_dCas9-TET1.dna

Additional file 10: Whole plasmid sequence for pMLM3636-Puro-dCas9-TET1-CGG.dna

Additional file 11: Whole plasmid sequence for pMLM3636-Puro-dCas9-TET1-PRM.dna

