## Supplementary table and figures for "MSH2 is not required for either maintenance of DNA methylation or repeat contraction at the *FMR1* locus in fragile X syndrome"

| ID # | Primer Name | Primer Sequence (5’-3’) |
|---|---|---|
| 1 | pX462-MSH2-dual-F | GAAAGGACGAAACACCGGTATGTGGATTCCATACAGTTAGAGCTAGAAATA |
| 2 | pX462-MSH2-dual-R | TTTCTAGCTCTAAAACCAGTTGATGGCCAGAGACAGCGGTGTTTCGTCCTTT |
| 3 | Gb_MSH2_Ex3F | TTTGGATTTTTCCTTTTTGCTT |
| 4 | Gb_MSH2_Ex3R | CCACATGCCTATACAAATGACA |
| 5 | *FMR1* exon1 F | GAACAGCGTTGATCACGTGAC |
| 6 | *FMR1* exon1 R | GTGAAACCGAAACGGAGCTGA |
| 7 | *GAPDH* exon1 F | TCGACAGTCAGCCGCATCT |
| 8 | *GAPDH* intron1 R | CTAGCCTCCCGGGTTTCTCT |
| 9 | FMR1-Met-IF | GGAATTTTAGAGAGGTYGAATTGGG |
| 10 | Gb_metbis-2645-R | aaacgacggccagtgCTCAAAAACTACCCTCCACC |
| 11 | Gb_FMR1-Met-2F | cacatcgctcagacacGTTATTGAGTGTATTTTTGTAGAAATGGG |
| 12 | Not_FraxC | AGTTCAGCGGCCGCGCTCAGCTCCGTTTCGGTTTCACTTCCGGT |
| 13 | FAM_Not_FraxR4 | CAAGTCGCGGCCGCCTTGTAGAAAGCGCCATTGGAGCCCCGCA |
| 14 | GAA-104F | GGCTTAAACTTCCCACACGTGTT |
| 15 | GAA-629R | AGGACCATCATGGCCACACTT |
| 16 | Gb_FXNMe-1212-F | cacatcgctcagacacGAGGTGAAATTTTTAGAGTTGTAGAATAGTTAGAGTAGTAG |
| 17 | Gb_FXNMe_1930-R | aaacgacggccagtgCACCTCCCAAAATACTAAAATTATAAACATAAACCA |
| 18 | PuroR-FMR1-gRNA-PRM_F | GAAAGGACGAAACACCgacagcgttgatcacgtgacggttttagagctaGAAAtagc |
| 19 | PuroR-FMR1-gRNA-PRM_R | TTTCtagctctaaaacggtcgaaagacagacgcgcgcGGTGTTTCGTCCTTT |
| 20 | Gb_3636-TET1-F | TCACTTTTTTTCAGGTTGGATACCCTCGTAAAGGCCACC |
| 21 | Gb_3636-TET1-R | TGTACTCGGTCATGGTGGCACCAGGGCCGGGATTCTCCTC |

| | | |
|---|---|---|
| 22 | Gb_dCas9-cmyc-F | CACTTCCTACCCTCGTAAAGGCCACCATGGGACCAGCCGCAAAGAGAGTG AAACTGGACGGAGGTCCTGCTGCAAAAAGGGTGAAGTTGGATGGAGACTAC AAAGACCATGACGGTG |
| 23 | gRNA-$CGG_6$ | acaccGGCGGCGGCGGCGGCGGCGGg |
| 24 | gRNA-PRM-1 | ACAGCGTTGATCACGTGACG |
| 25 | gRNA-PRM-2 | CGCGCGTCTGTCTTTCGACC |
| 26 | gRNA-MSH2-1 | GGTATGTGGATTCCATACAG |
| 27 | gRNA-MSH2-2 | CAGTTGATGGCCAGAGACAG |
| 28 | 3636-727F | AAGCAGGCTTTAAAGGAACCA |
| 29 | 459-377R | TACCCGTTACATAACTTACG |

**Table S1**: Primer and Guide RNA Sequences

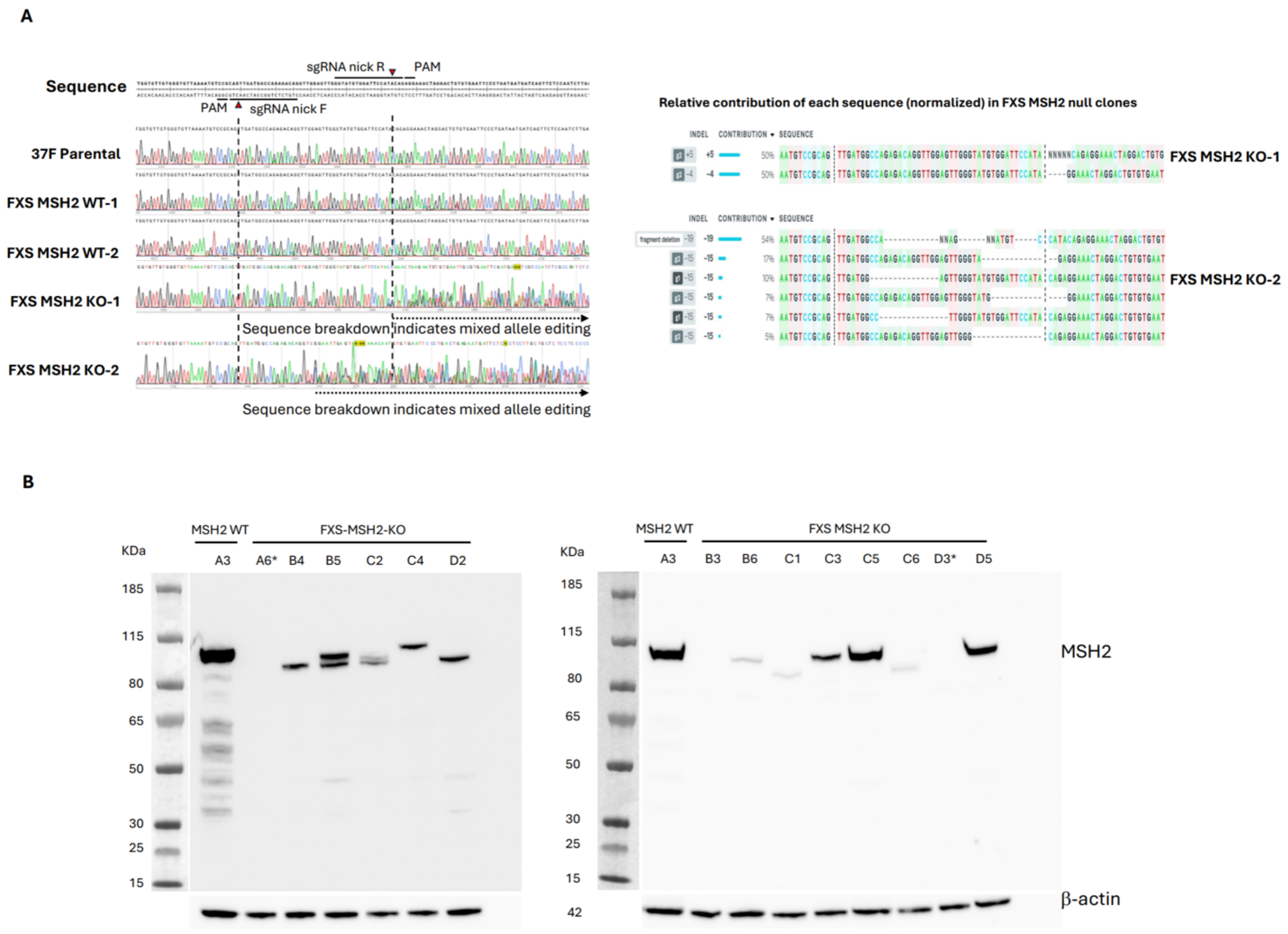


**Figure S1. (A)** Sanger Sequencing results for MSH2 exon 3 PCR on FXS MSH2 WT and FXS MSH2 KO cell lines aligned to MSH2 exon 3. Diagram shows location of nickase CRISPR guide RNAs and single strand break locations. FXS MSH2 WT samples show 100% match to reference sequence and both FXS MSH2 KO samples display sequence breakdown within the two CRISPR nick sites indicative of CRISPR editing and mixed allele population. Sequence analysis using Synthego ICE software shows complete loss of *MSH2* exon 3 allele in both cell lines, with indel regions that match the region of sequence breakdown. **(B)** Western blot for MSH2 with b-actin loading control on single-cell clones of WCMC 37F FXS ESCs edited with dual CRISPR nickase targeting *MSH2* exon 3. All clones were analyzed here, and complete knockouts A6 and D3 (marked with asterisks) were selected for further study and renamed FXS MSH2 KO-1 and FXS MSH2 KO-2.

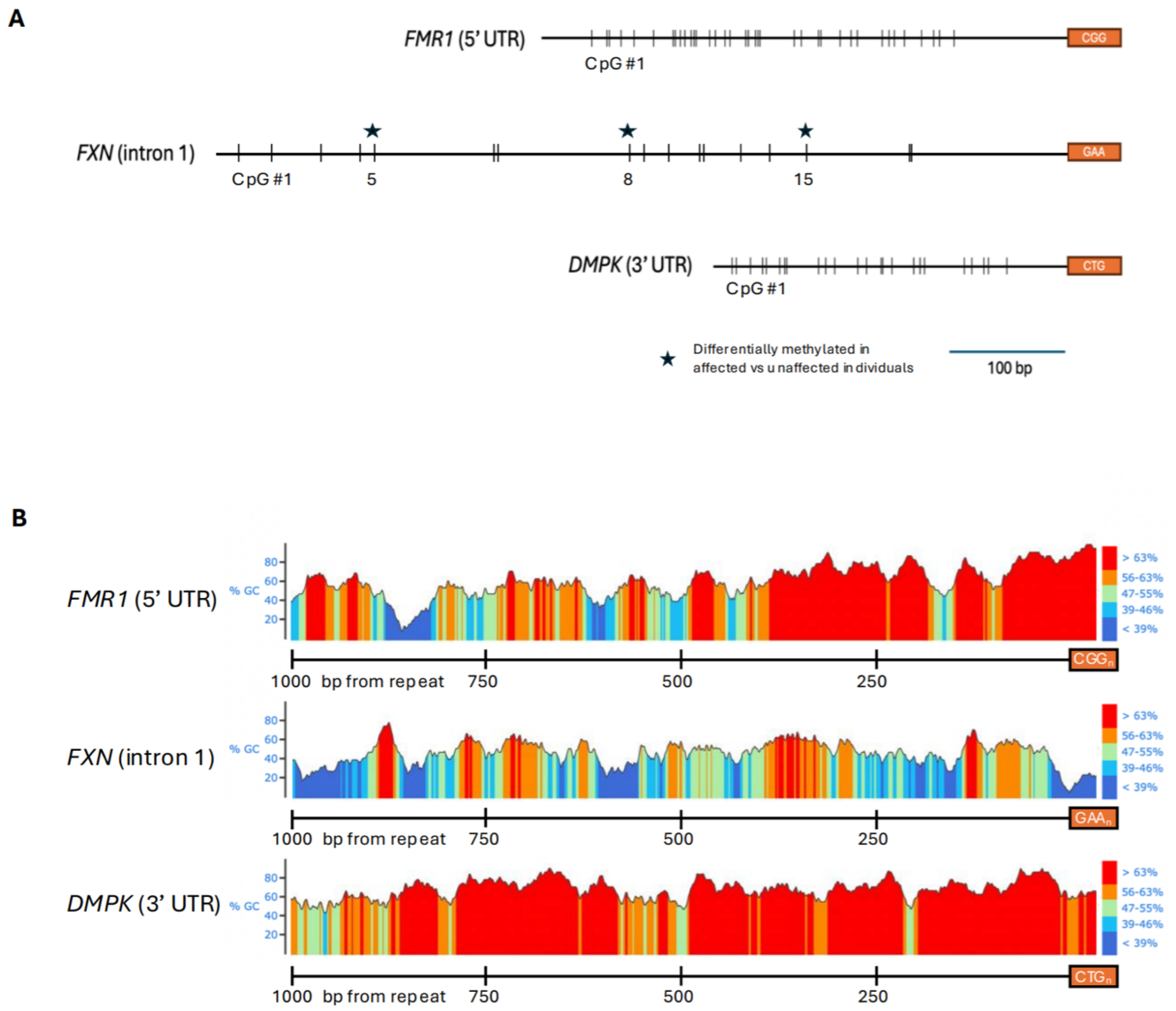


**Figure S2. (A)** Diagram showing CpG residues that are methylated in regions upstream of the disease-causing repeats in Fragile X syndrome (*FMR1),* Friedreich's Ataxia (*FXN),* and Myotonic Dystrophy Type I (*DMPK).* CpGs that are differentially methylated in FRDA vs non-FRDA lymphoblastoid cell lines (23) are noted with an asterisk and numbered. **(B)** Overall GC content of the region 1 kb upstream of the repeats in each disorder. The height of the curve and the color of the area under the curve are based on the average GC content (%) from low (blue) to high (red).

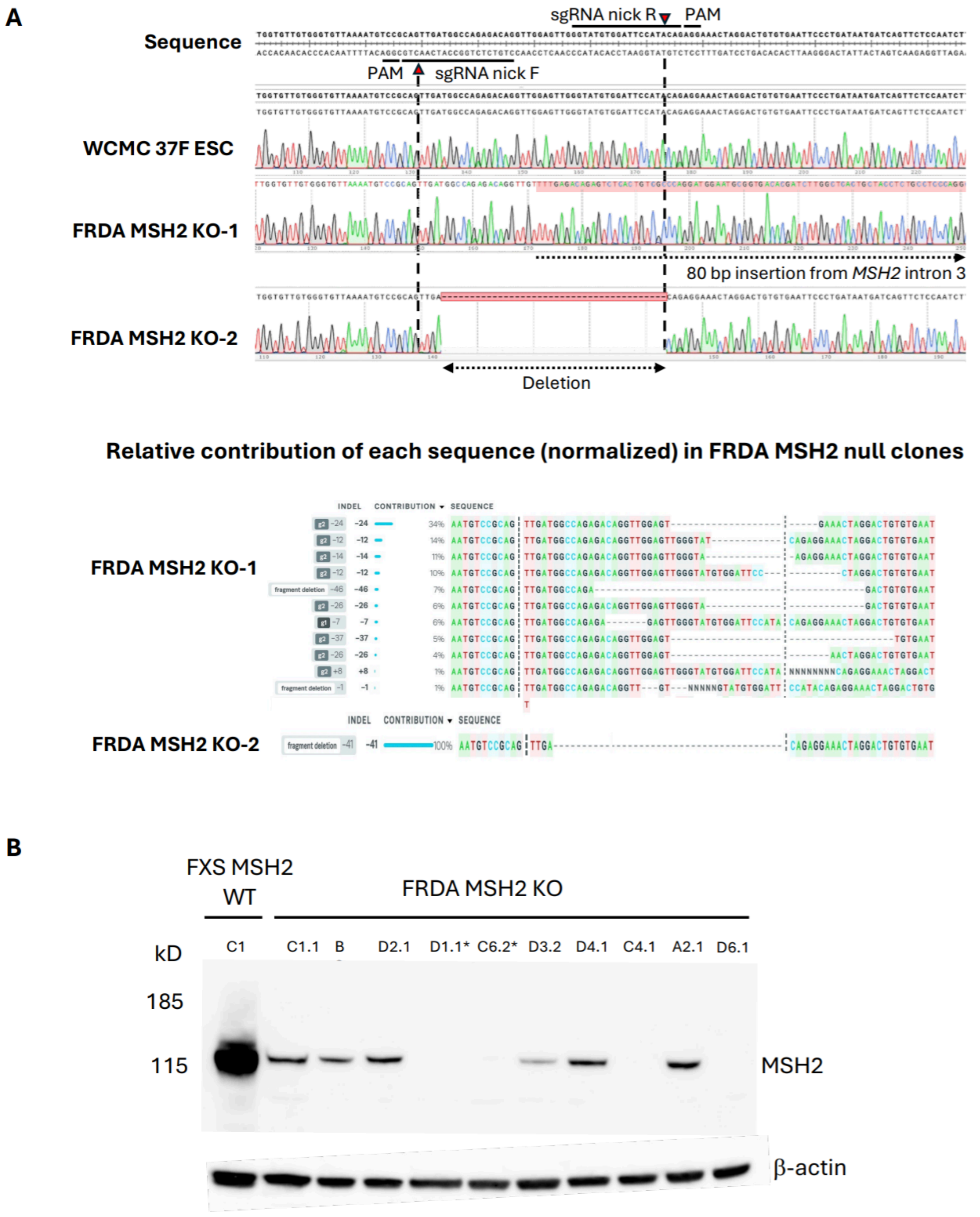


**Figure S3. (A)** Sanger Sequencing results for MSH2 exon 3 PCR on FRDA MSH2 KO cell lines aligned to MSH2 exon 3. Diagram shows location of nickase CRISPR guide RNAs and single strand break locations. Both FRDA MSH2 KO samples show evidence of CRISPR editing:

FRDA MSH2 KO-1 has an 80 bp insertion of a region of *MSH2* intron 3 inserted in between the CRISPR cut sites and FRDA MSH2 KO-2 has a 41 bp deletion with the 3' cut site matching the sgRNA nick R cut site. Sequence analysis using Synthego ICE software shows complete loss of *MSH2* exon 3 allele in both cell lines, with a deletion matching the Sanger sequencing alignment in FRDA MSH2 KO-2. In FRDA MSH2 KO-1, ICE did not identify the insertion, but it did confirm complete loss of the typical *MSH2* allele. **(B)** Western blot for MSH2 with β-actin loading control on single-cell clones of GM23404 FRDA human iPSCs edited with dual CRISPR nickase targeting *MSH2* exon 3. All clones isolated are analyzed here and 4 clones were found to be complete knockouts: D1.1, C6.2, C4.1, D6.1. We selected two clones C6.2 and D1.1 for continued study (marked with asterisk in the Figure) and renamed them FRDA MSH2 KO-1 and FRDA MSH2 KO-2.

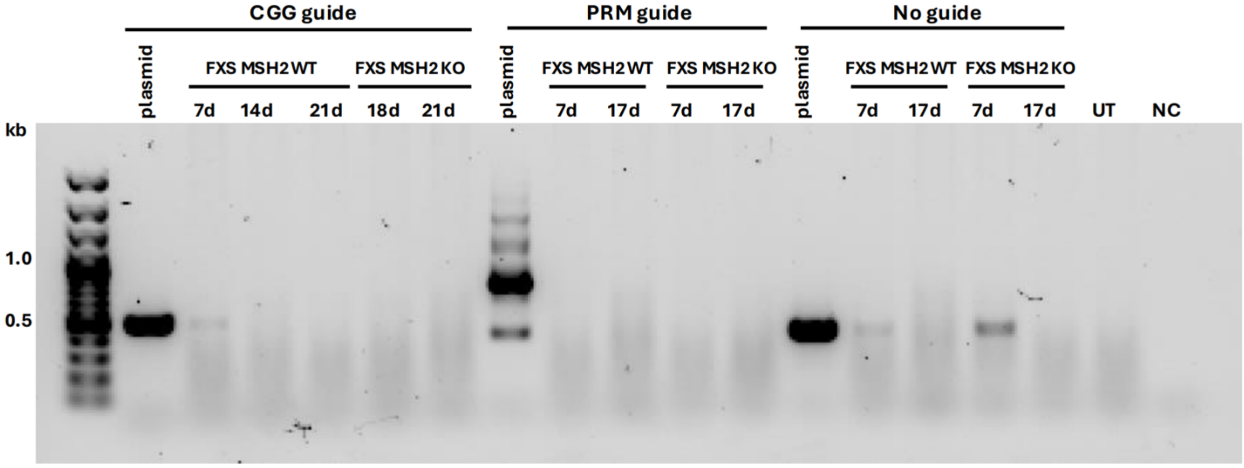


**Figure S4**. PCR showing that low levels of TET1 plasmids are present at day 7 after transfection and completely lost by 2 weeks post transfection.

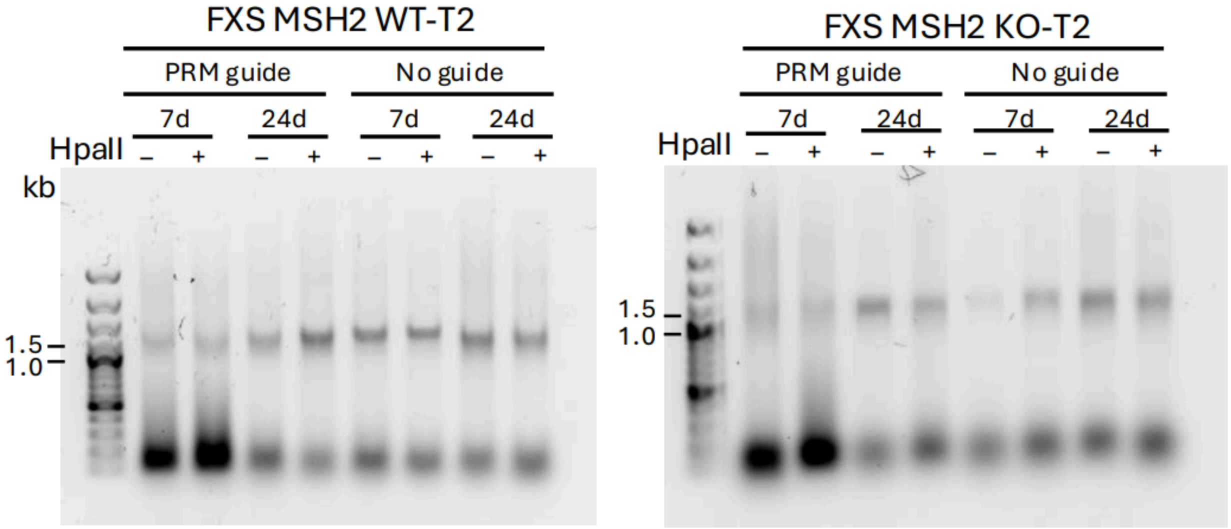


**Figure S5**. CGG repeat PCR results for DNA from replicate transfection (T2) at 7d and 24d for FXS MSH2 WT-2 and FXS MSH2 KO-2 cells transfected with plasmid dCas9-TET1-PRM and plasmid dCas9-TET1 without any gRNAs. HpaII-digested DNA lanes show amplification of alleles that are fully methylated while undigested samples amplify both methylated and unmethylated alleles. The *FMR1* gene in FXS ESCs carries 400 repeats resulting in a PCR product of about 1.4 kb. FXS MSH2 WT and FXS MSH2 KO cell lines transfected with dCas9-TET1-PRM and dCas9-TET1 with no guide show CGG repeat sizes similar to untransfected (UT) cells at day 7 (7d) and day 24 (24d). The results of CGG repeat PCR for transfection T1 are shown in Figure 7C.
