## additional file 7 for "MSH2 is not required for either maintenance of DNA methylation or repeat contraction at the *FMR1* locus in fragile X syndrome": methylation analysis script.py

```
filename = input("Enter the name of the alignment file: ")
data = open(filename, 'r')
alignDict = {}
nameDict = {}
for line in data:
    # collect the alignments
    if line[:5] == "Query":
        field = line.strip().split()
        if field[0] in alignDict:
            alignDict[field[0]] = alignDict[field[0]] + field[2]
        else:
            alignDict[field[0]] = field[2]
    # map the queryID to local ID
    elif line[:7] == "Subject":
        field = line.strip().split()
        name = field[1].split(":")
        qId = field[3].split("|")
        nameDict[qId[1]] = name[1]
    else: pass
data.close()

# find the position of all the Cs in CpGs
alignto = alignDict["Query"].upper()
del alignDict["Query"]
isC = []
lenQuery = len(alignto)
for i in range(len(alignto)):
    if alignto[i] == "C":
        isC.append(i)
    else: pass

# set up lists to later calculate % methylation at each position
Cpcnt = [0] * len(isC)
tpcnt = [0] * len(isC)

# count the number of alleles with a given % of methylation in bins of 10
allPcnt = [0] * 10

# count the number of unique alleles
allType = {} 

# then see if those positions are C or T in the test sequences
for id in sorted(alignDict):
    checkC = alignDict[id]
    # equalise the lengths of the test sequences
    if len(checkC) != lenQuery:
        checkC = checkC + ((lenQuery - len(checkC)) * "X")
    else: pass
    # collect the C or t value and update % methylation at each position
    cCount = ""
    for i, posn in enumerate(isC):
        if checkC[posn] == ".":
            cCount += "C"
            Cpcnt[i] +=1
        elif checkC[posn] == "T":
            cCount += "t"
            tpcnt[i] +=1
        else:
            cCount += "-"
    # get the pcent methylation
    numC = cCount.count("C")
    numt = cCount.count("t")
    pcent = numC/(numC + numt)
    pcentIndex = int(10 * pcent)
    if pcentIndex == 10:
        pcentIndex = 9
    else: pass
    allPcnt[pcentIndex] +=1
    # check for allele uniqueness
    if cCount in allType:
        allType[cCount] +=1
    else:
        allType[cCount] =1

# calculate % methylation at each position and prepare output
cpgNum = ""
cpgVal = ""
for i in range(len(Cpcnt)):
    cpgNum += str(i + 1) + "\t"
    cpgVal += str(round(Cpcnt[i]/(Cpcnt[i] + tpcnt[i]), 2)) + "\t"

outfile = open("Methylation_statistics.txt", 'w')
print("Number of alleles: ", len(alignDict), file=outfile)
print("", file=outfile)
print("Percent methylation of each CpG", file=outfile)
print(cpgNum, file=outfile)
print(cpgVal, file=outfile)
print("", file=outfile)
print("Number of alleles with methylation percent", file=outfile)
print("<10\t<20\t<30\t<40\t<50\t<60\t<70\t<80\t<90\t90+", file=outfile)
methpcent = ""
for value in allPcnt:
    methpcent += str(value) + "\t"
print(methpcent, file=outfile)
print("", file=outfile)
print("Unique alleles found", file=outfile)
print("Frequency\tNumt\tNumdash\tNumC\tSequence", file=outfile)
sortAllele = sorted(allType.items(), key=lambda x:x[1], reverse=True)
for allele in sortAllele:
    outLine = str(allele[1]) + "\t" + str(allele[0].count("t")) + "\t" + str(allele[0].count("-")) + "\t" + str(allele[0].count("C"))
    for char in allele[0]:
        outLine += "\t" + char
    print(outLine, file=outfile)
outfile.close()
```
